# A new type of mouse gaze shift is led by directed saccades

**DOI:** 10.1101/2021.02.10.430669

**Authors:** Sebastian H. Zahler, David E. Taylor, Joey Y. Wong, Julia M. Adams, Evan H. Feinberg

**Author notes:** These authors contributed equally (Sebastian H. Zahler, David E. Taylor).

## Abstract

Animals investigate their environments by directing their gaze towards salient stimuli. In the prevailing view, mouse gaze shifts are led by head rotations that trigger compensatory, brainstem-mediated eye movements, including saccades to reset the eyes. These “recentering” saccades are attributed to head movement-related vestibular and optokinetic cues. However, microstimulating mouse superior colliculus (SC) elicits directed head and eye movements that resemble SC-dependent sensory-guided gaze shifts made by other species, raising the possibility mice generate additional types of gaze shifts. We investigated this possibility by tracking eye and attempted head movements in a head-fixed preparation that eliminates head movement-related sensory cues. We found tactile stimuli evoke gaze shifts involving directed saccades that precede attempted head rotations. Optogenetic perturbations revealed SC drives touch-evoked gaze shifts. Thus, mice make sensory-guided, SC-dependent gaze shifts led by directed saccades. Our findings uncover diversity in mouse gaze shifts and provide a foundation for studying head-eye coupling.

## Introduction

Natural environments are complex and dynamic, and animals frequently redirect their gaze to scrutinize salient sensory stimuli. Gaze shifts employ head and eye movement coupling strategies that depend on context and can vary between species (Goldring et al., 1996; Land, 2019; Land and Nilsson, 2012; Populin, 2006; Populin and Rajala, 2011; Populin et al., 2004a; Ruhland et al., 2013; Tollin et al., 2005). Mice are an increasingly important model organism in vision research, yet the strategies they use to shift their gaze remain incompletely understood. Revealing these strategies is essential to understanding mouse visual ethology and the underlying neural mechanisms.

The prevailing view holds that species such as mice whose retinae lack high-acuity specializations (afoveates) generate gaze shifts driven by head movements followed by “recentering” saccades (Land and Nilsson, 2012). Indeed, recent studies tracking head and eye movements in freely moving mice found that spontaneous and visually evoked mouse gaze shifts matched this description (Meyer et al., 2018, 2020; Michaiel et al., 2020; Payne and Raymond, 2017). Specifically, during a gaze shift, slow eye movements stabilize the retinal image by countering the rotation of the head and are punctuated by fast saccadic eye movements to recenter the eyes in the orbits as they approach the end of their range of motion. These recentering saccades—also known as “compensatory” saccades or the quick phase of nystagmus—are centripetal, occur in the direction of the head movement, and are thought to be driven by vestibular or optokinetic signals acting on circuits in brainstem (Curthoys, 2002; Hepp et al., 1993; Kitama et al., 1995; Meyer et al., 2020; Michaiel et al., 2020; Payne and Raymond, 2017). These recent observations have buttressed the view that gaze shifts in mice and other afoveates are led by head movements, with eye movements made only to compensate for the effects of head movements. In contrast, primates and other foveate species are capable of an additional form of gaze shift led by directed saccades, with or without directed head movements, to redirect their gaze towards salient stimuli (Bizzi et al., 1972; Freedman, 2008; Lee, 1999; Zangemeister and Stark, 1982). Directed saccades differ from recentering saccades in that they have endpoints specified by the location of the stimulus (and therefore are often centrifugally directed), typically occur simultaneously with or slightly before (20-40 ms) head movements during gaze shifts, and are driven by midbrain circuits, particularly the superior colliculus (SC). To date, there is no behavioral evidence that mice or any afoveate species generate directed saccades or gaze shifts led by eye movements.

However, three observations are inconsistent with the model that all saccades in mice are exclusively recentering and made to compensate for head movements. First, mouse saccades are not only a product of vestibular or optokinetic cues, because head-fixed mice, in which these signals do not occur, generate saccades, albeit less frequently. Second, neuroanatomical and functional studies suggest that the circuits that underlie directed saccades are conserved in mice (May, 2006; Sparks, 1986, 2002). Specifically, microstimulation of the mouse superior colliculus (SC) showed that it contains a topographic map of saccade and head movement direction and amplitude (Masullo et al., 2019; Wang et al., 2015) roughly aligned with maps of visual, auditory, and somatosensory space (Drager and Hubel, 1975, 1976). These SC sensory and motor maps resemble those believed to underlie primates’ and cats’ ability to make gaze shifts led by directed saccades towards stimuli of these modalities (Sparks, 1986, 2002). Third, saccade-like eye movements occurring in the absence of head movements have occasionally been observed in freely moving mice, albeit infrequently and usually in close proximity to head movements (Meyer et al., 2018, 2020; Michaiel et al., 2020).

We therefore hypothesized that mice innately generate gaze shifts that incorporate directed saccades. We predicted that this ability was obscured in previous studies for several reasons. First, in freely moving mice it is difficult to uncouple the contributions of reafferent vestibular and optokinetic inputs from those of exafferent (extrinsic) sensory inputs to saccade generation. Second, previous analyses in mice were mostly confined to spontaneous or visually guided gaze shifts, and there is evidence in humans, non-human primates, and cats that gaze shifts in response to different sensory modalities can involve distinct head-eye coupling strategies (Goldring et al., 1996; Populin, 2006; Populin and Rajala, 2011; Populin et al., 2004a; Ruhland et al., 2013; Tollin et al., 2005). Third, in freely moving mice it is difficult to present stimuli in specific craniotopic locations. We therefore reasoned that by using a head-fixed preparation both to eliminate vestibular and optokinetic cues and to present stimuli of different modalities at precise craniotopic locations, we could systematically determine whether mice are capable of gaze shifts involving saccades whose endpoints depend on stimulus location and show different coupling to head movements. We found that tactile stimuli evoke saccades whose endpoints depend on stimulus location, that these saccades precede attempted head rotations, and that these touch-evoked gaze shifts are SC-dependent. Together, these results resolve an apparent discrepancy between mouse neuroanatomy and behavior, demonstrating that mice are capable of generating gaze shifts led by directed saccades.

## Results

### Stimulus-evoked gaze shifts in head-restrained mice

To test the hypothesis that mice possess an innate ability to make sensory-evoked gaze shifts that incorporate directed saccades coupled to head movements, we head-fixed naïve, wild-type adult animals and used infrared cameras to track both pupils and a strain gauge (also known as a load cell) to measure attempted head rotations. Previous studies in head-fixed mice observed occasional undirected saccades in response to changes in the visual environment (Samonds et al., 2018) and visually guided saccades only after weeks of training and at long (∼1 s) latencies (Itokazu et al., 2018). We therefore tested a panel of stimuli of different modalities to determine whether they could evoke saccades. We began by testing the following stimuli from a constant azimuthal location: 1) a multisensory airpuff that provides tactile input to the ears and generates a loud, broadband sound; 2) an auditory stimulus consisting of the same airpuff moved away from the animal so as not to provide tactile input; 3) a tactile stimulus consisting of a bar that nearly silently taps the ear; and 4) a visual stimulus consisting of a bright LED. Stimuli were delivered on either side of the animal every 7-12 s in a pseudorandom sequence (Fig. 1A). The probability of horizontal eye movements increased sharply and significantly above the low baseline level (1.3 ± 0.2%, mean ± s.d.) in the 100 ms period following delivery of multisensory airpuffs (ear airpuff: 29.0 ± 7.5%, p < 0.001 paired Student’s t-test; whisker airpuff: 12.5 ± 2.3%, p < 0.001), auditory airpuffs (3.5 ± 1.2%, p < 0.05), and tactile stimuli (4.5 ± 0.5%, p < 0.001) and remained slightly elevated for at least 500 ms (Fig. 1C-F, H-K; Supplementary Figure 2). In contrast, the probability of saccade generation was not changed by visual stimuli (Fig. 1G, L; 1.3 ± 0.2%, p = 0.61). We consider these stimulus-evoked eye movements to be saccades because they reached velocities of several hundred degrees per second (Supplementary Fig. 1), displayed a main sequence, *i.e.*, peak velocity scaled linearly with amplitude (Supplementary Fig. 1), and were bilaterally conjugate (Supplementary Fig. 1) (Bahill et al., 1975). As in previous studies, saccade size for temporal-to-nasal movements was slightly larger than for nasal-to-temporal movements (Meyer et al., 2018), but this asymmetry was eliminated by averaging the positions of both pupils (before averaging: temporal saccade amplitude = 10.9 ± 0.8° (mean ± s.d.), nasal saccade amplitude = 8.4 ± 1.1°, p = 0.0009; after averaging, leftward amplitude = 10.2 ± 1.0°, rightward amplitude = 9.3 ± 0.4°, p = 0.0986). We next examined attempted head movements. The baseline frequency of attempted head movements was much higher than that of eye movements (27.8 ± 4.8%, mean ± s.d.). Mirroring results for saccades, auditory, tactile, and audiotactile stimuli evoked attempted head movements but visual stimuli did not (Fig. 1M-Q; auditory: 53.2 ± 21.9%, p = 0.045; tactile: 67.1 ± 10%, p < 0.01; ear airpuff: 86.6 ± 8.7%, p < 10^-5^; whisker airpuff: 79.2 ± 9.5%, p < 0.001; visual: 26.9 ± 2.0%, p = 0.0505, paired Student’s t-test). These data demonstrate that both auditory and tactile stimuli are sufficient to evoke gaze shifts in head-fixed mice, and that mice make sensory-evoked saccades in the absence of vestibular and optokinetic inputs.

**Figure 1.**
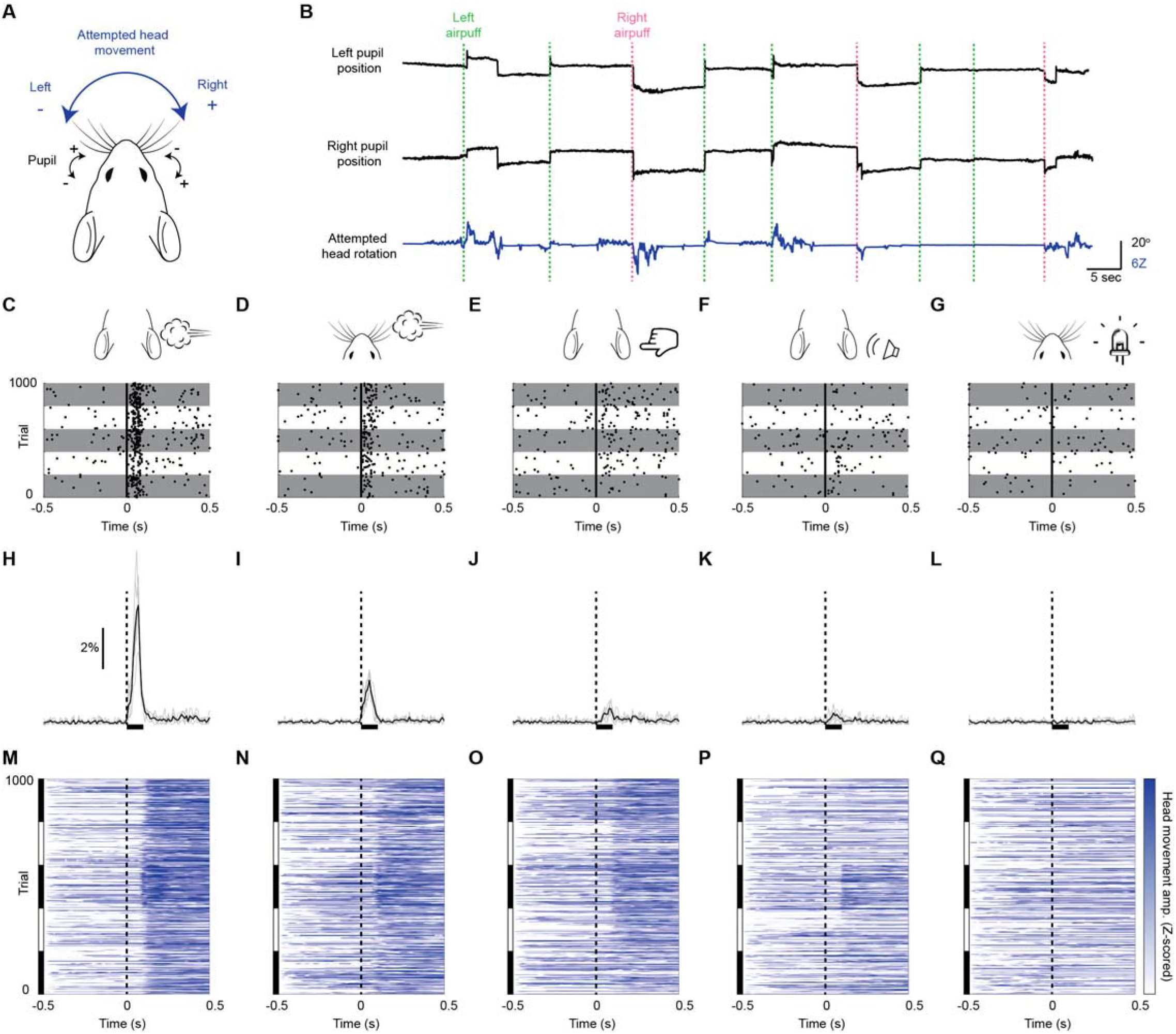
Mice innately make sound- and touch-evoked gaze shifts. **(A)** Behavioral schematic. Naïve mice are head-fixed and stimuli are presented on either side. Both eyes are tracked using cameras and attempted head rotations are measured using a strain gauge (load cell). In subsequent quantification, eye positions to the right of center (nasal for left eye, temporal for right eye) are positive, and eye positions to the left of center (temporal for left eye, nasal for right eye) are negative, with zero defined as the mean eye position. Likewise, attempted rightward head movements are positive, and leftward head movements are negative. **(B)** Sample eye and attempted head movement traces. Dashed vertical lines indicate left (magenta) and right (green) ear airpuff delivery. **(C-G)** Saccade rasters for 5 representative mice in response to (C) ear airpuffs, (D) whisker airpuffs, (E) ear tactile stimuli, (F) auditory airpuffs, and (G) visual stimuli. Each row corresponds to a trial. Each dot indicates onset of a saccade. Vertical black lines denote time of left or right stimulus delivery. Each gray or white horizontal stripe contains data for a different mouse. n = 1000 randomly selected trials (200/mouse). **(H-L)** Peri-stimulus time histograms showing instantaneous saccade probabilities in response to (H) ear airpuffs, (I) whisker airpuffs, (J) ear tactile stimuli, (K) auditory airpuffs, and (L) visual stimuli for mice from (C-G). Each light trace denotes a single animal; black traces denote population mean. Dashed lines denote time of stimulus delivery. Horizontal bar indicates the 100 ms response window used in subsequent analyses. **(M-N)** Heatmaps of attempted head movements in response to (M) ear airpuffs, (N) whisker airpuffs, (O) ear tactile stimuli, (P) auditory airpuffs, and (Q) visual stimuli for mice from (C-L). Each row corresponds to an individual trial from C-G. Black and white bars at left indicate blocks of trials corresponding to each of 5 different mice. Dashed line denotes stimulus delivery time.

### Tactile stimuli evoke directed saccades whose endpoints depend on stimulus location

To determine whether sensory-evoked saccades are directed, we asked whether saccade endpoints are dependent on stimulus location. We began by examining the endpoints of saccades evoked by left and right ear airpuffs (Fig. 2A). We found that left ear airpuffs evoked saccades with endpoints far left of center (with center defined as the mean eye position), whereas right ear airpuffs evoked saccades with endpoints far right of center (left: −5.4 ± 4.5°, right: 5.4 ± 3.4°, mean ± s.d., p < 10^-10^ Welch’s t-test, n = 2155 trials). To understand how this endpoint segregation arises, we examined the trajectories of individual saccades (Fig. 2E). We found that left ear airpuffs elicited nearly exclusively leftward saccades (94.2 ± 3.1%, mean ± s.d., n = 5 mice), whereas right ear airpuffs elicited nearly exclusively rightward saccades (96.0 ± 2.1%)—often from the same eye positions. By definition, from any eye position, one of these directions must lead away from center and is thus centrifugal rather than centripetal. In addition, puff-evoked saccades that began towards the center often reached endpoints at eccentricities of 5 to 10 degrees. To further test whether saccade endpoints are specified by stimulus location, we repositioned the airpuff nozzles to stimulate the whiskers and repeated the experiments. We reasoned that saccade endpoints should become less eccentric as stimulus eccentricity decreases. Indeed, airpuffs applied to the whiskers evoked saccades with endpoints central to those evoked by ear airpuff stimulation, such that the ordering of saccade endpoints mirrored that of stimulus locations (Mean endpoints: left ear, −5.4 ± 4.5°; left whiskers, −0.4 ± 3.8°; right whiskers, 0.8 ± 3.9°; right ear, 5.4 ± 3.4°; mean ± s.d., p < 0.05 for all pairwise comparisons, paired two-tailed Student’s t-test) (Fig. 2A,B; Supplementary Fig. 3). In this cohort, the separation between whisker-evoked saccade endpoints, although significant, was small, but in other cohorts we have observed larger separation (as well as higher auditory-evoked saccade probabilities) (Supplementary Fig. 4). Taken together, these data suggest that touch-evoked saccades are directed towards particular eye positions that are specified by stimulus location.

**Figure 2.**
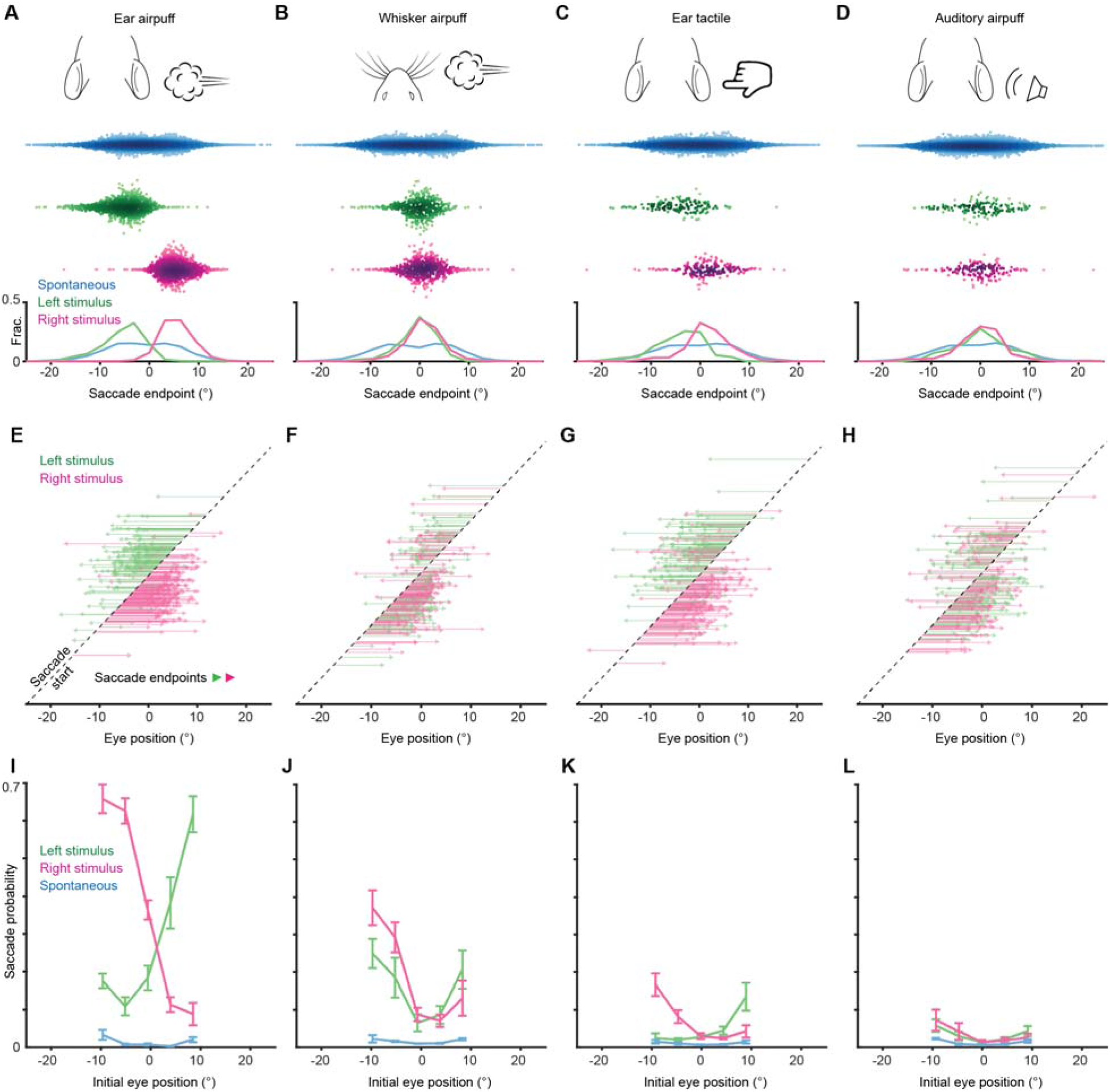
Endpoints and trajectories of sensory-evoked saccades. **(A-D)** Endpoints for ear airpuff-, whisker airpuff-, ear tactile-, and auditory airpuff-evoked saccade. Top, schematics of stimuli. Middle, scatter plots showing endpoints of all saccades for all animals (n=see below, 5 animals) made spontaneously (blue) and in response to left (green) and right (magenta) stimuli. Darker shading indicates areas of higher density. Bottom, histograms of endpoint distributions for spontaneous and evoked saccades. **(E-H)** Trajectories of individual stimulus-evoked saccades. Each arrow denotes the trajectory of a single saccade. Saccades are sorted according to initial eye positions, which fall on the dashed diagonal line. Saccade endpoints are indicated by arrowheads. Because the probability of evoked gaze shifts differed across stimuli, data for ear and whisker airpuffs are randomly subsampled (15% and 30% of total trials, respectively) to show roughly equal numbers of trials for each condition. **(I-L)** Relationship between eye position and saccade probability. Green and magenta lines indicate population means (n = 5 mice) for saccades evoked by left and right stimuli, respectively. Blue lines indicate spontaneous saccades. Error bars indicate s.e.m. Saccade numbers in A-H: ear airpuff sessions, spontaneous = 7146, left ear airpuff-evoked = 942 (141 in E), right ear airpuff-evoked = 1213 (182 in E); whisker airpuff sessions: spontaneous = 7790, left whisker airpuff-evoked = 440 (132 in F), right whisker airpuff-evoked = 606 (181 in F); ear tactile sessions, spontaneous = 6706, left ear tactile-evoked = 133, right ear tactile-evoked = 186; auditory sessions, spontaneous = 10240, left auditory-evoked = 140, right auditory-evoked = 158.

We next examined the endpoints of saccades evoked by tactile and auditory stimuli. Similar to the multisensory airpuff stimuli, tactile stimuli delivered to the left and right ears evoked saccades whose endpoints were significantly different (−3.9 ± 5.3° (left) vs. 1.5 ± 5.0° (right); mean ± s.d., p < 10^-10^, Welch’s t-test, n = 452 trials) and whose directions were largely opposite from nearly all eye positions (left stimuli evoked 77.5 ± 23.7% leftward saccades, right stimuli evoked 78.3 ± 11.6% rightward saccades, n = 5 mice). This result suggests that tactile stimuli are sufficient to induce gaze shifts that involve directed saccades. We next examined the endpoints and trajectories of saccades evoked by left and right auditory stimuli (Fig. 2D, H). Strikingly, the saccade endpoint locations did not differ significantly for left and right auditory stimuli and were located centrally (0.1 ± 5.1° (left) vs. −0.1 ± 5.1° (right); mean ± s.d., p = 0.72, Welch’s t-test, n = 298 trials). Because we had fewer trials with sound-evoked gaze shifts overall, to confirm that this lack of statistically significant endpoint separation was not a result of lower statistical power, we repeated our analyses on matched numbers of sound- and touch-evoked saccades sampled at random, once again observing that left and right ear airpuff-, whisker airpuff-, and ear tactile-evoked saccade endpoints were significantly different (ear airpuff, p < 10^-10^; whisker airpuff, p = 0.0039; ear tactile, p < 10^-10^; auditory airpuff, p = 0.50; Welch’s t-test, see figure for complete statistics). The central endpoints of sound-evoked saccades arose because, in contrast to touch-evoked saccades, saccades evoked by both right and left stimuli traveled in the same, centripetal direction from all initial eye positions: rightward from eye positions to the left of center, and leftward from eye positions to the right of center (fraction centripetal: left airpuff for initial eye positions left of center, 0.89 ± 0.08; left airpuff for initial eye positions right of center, 0.90 ± 0.09; right airpuff for initial eye positions left of center, 0.87 ± 0.07; right airpuff with initial eye positions right of center, 0.76 ± 0.12).. We compared the mean sound-evoked saccade endpoint location to the mean overall eye position and found no significant difference, suggesting that auditory saccades function to recenter (p = 0.93, one-sample Student’s t-test) (Land and Nilsson, 2012; Meyer et al., 2018, 2020; Michaiel et al., 2020; Paré and Munoz, 2001; Tatler, 2007). Thus, both the auditory and tactile components of the airpuff stimuli are sufficient to evoke saccades but only tactile stimulation evokes directed saccades whose endpoints are specified by the site of stimulation.

### Starting eye position determines saccade probability

We observed that gaze shifts were generated on a subset of trials. We therefore sought to understand factors underlying the probabilistic nature of evoked gaze shifts. We first examined arousal, which modulates many neuronal and behavioral phenomena, by inferring arousal levels from pupil diameter. Surprisingly, saccade probability did not differ between trials with high (dilated pupils) or low (constricted pupils) arousal (Supplementary Fig. 5) (Reimer et al., 2014). Next, we asked whether novelty plays a role by examining the effects of sensory history. Saccade probability was constant within sessions but declined somewhat over successive sessions, suggesting that sensory history plays a role on long time scales (Supplementary Fig. 5). We then examined starting eye position. Strikingly, saccade probability varied strongly with initial eye position (Fig. 2I-L, Supplementary Fig. 4I-L). In addition, the relationship between initial eye position and saccade probability differed across stimuli, with the lowest probability coinciding with the mean endpoint of saccades evoked by that stimulus (Fig. 2A-D, I-L).

### Stimulus-evoked gaze shifts involve a different type of head-eye coupling

Next, we examined the relationship between attempted head rotations and saccades during gaze shifts. We first analyzed the relative timing of the head and eye components of spontaneous gaze shifts. Previous studies in freely moving and head-fixed mice found that on average, spontaneous and visually evoked saccades in freely moving mice are preceded by head rotations, and that spontaneous saccades in head-fixed mice are similarly preceded by attempted head rotations (Meyer et al., 2018, 2020; Michaiel et al., 2020; Payne and Raymond, 2017). Consistent with these observations, we found that on average spontaneous saccades were preceded by attempted head rotations (Fig. 3B, E). Interestingly, attempted head rotations during spontaneous saccades appeared biphasic, with a slower phase beginning before saccades (median: 90 ms before saccade onset) followed by a fast phase beginning slightly after saccade onset. This biphasic response closely resembles the eye-head coupling pattern reported in freely moving mice (Meyer et al., 2020).

**Figure 3.**
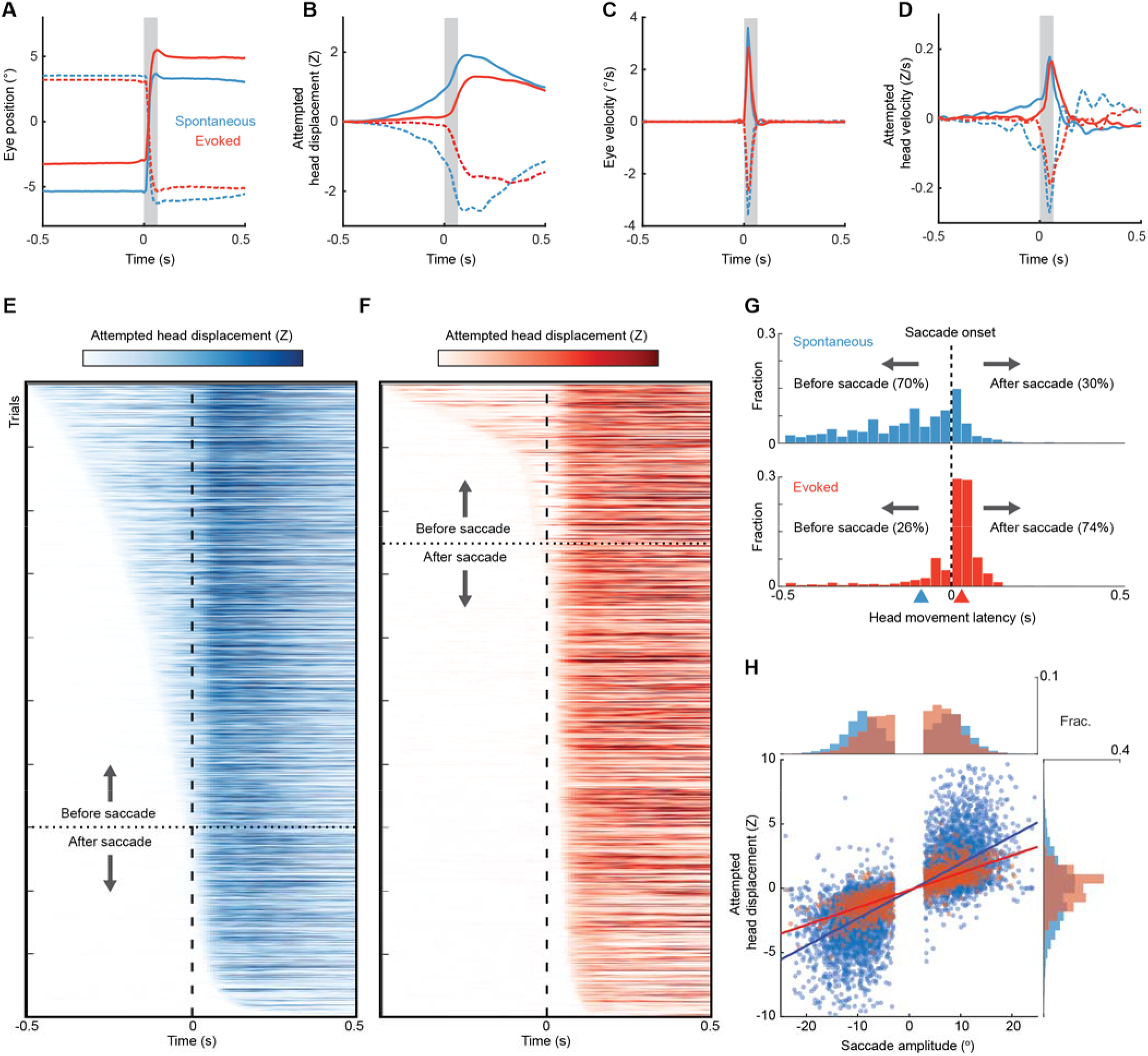
Different head-eye coupling during spontaneous and touch-evoked gaze shifts. **(A)** Mean trajectories of all rightward (solid traces) and leftward (dashed traces) saccades during spontaneous (blue, n= 7146) and ear airpuff-evoked (red, n = 1437) gaze shifts. Means + s.e.m. (smaller than line width). Gray bar indicates average saccade duration. **(B)** Mean attempted head movement amplitudes accompanying rightward (solid traces) and leftward (dashed traces) saccades during spontaneous (blue) and ear airpuff-evoked (red) gaze shifts. **(C)** Mean velocities of all rightward (solid traces) and leftward (dashed traces) saccades during spontaneous (blue) and ear airpuff-evoked (red) gaze shifts. (**D**) Mean attempted head movement velocities accompanying rightward (solid traces) and leftward (dashed traces) saccades during spontaneous (blue) and ear airpuff-evoked (red) gaze shifts. **(E, F)** Timing of attempted head movements relative to saccades during all spontaneous (E) and ear airpuff-evoked (F) gaze shifts. Each row corresponds to a single gaze shift. Darker shades indicate larger attempted head displacement. Dashed vertical line indicates time of saccade onset. Trials are sorted according to latency of attempted head movements. Trials above and below dashed horizontal line correspond to attempted head movements that began before and after saccades, respectively. **(G)** Distributions of attempted head movement latencies relative to saccade onset for spontaneous (top) and ear airpuff-evoked (bottom) saccades. Medians indicated by blue and red arrowheads. Distributions are significantly different (p < 10^-5^, permutation test). **(H)** Head-eye amplitude coupling of spontaneous (blue) and ear airpuff-evoked saccades (red). Each dot corresponds to a single gaze shift. Attempted head amplitude was measured 150 ms after saccade onset. Spontaneous: R^2^ = 0.58, slope = 0.214, p < 10^-10^. Evoked: R^2^ = 0.69, slope = 0.135, p < 10^-10^. Spontaneous and evoked regression slopes were significantly different (p = 0.01, permutation test). Histograms above and beside scatter plot indicate distributions of saccade and attempted head movement amplitudes, respectively. Mean head and eye movement amplitudes were significantly smaller for airpuff-evoked saccades (p < 10^-5^, permutation test).

We next asked whether the relative timing of eye and attempted head movements was similar for stimulus-evoked gaze shifts. Surprisingly, ear airpuff-evoked saccades were not preceded by slow attempted head rotations but were nevertheless accompanied by fast attempted head rotations that began slightly after saccade onset (Fig. 3B). This pattern was mirrored in the average eye and attempted head movement traces for whisker airpuff-, auditory airpuff-, and ear tactile-evoked saccades (Supplementary Fig. 6A-D). To better understand how these patterns arose, we examined head and eye movement timing at the single-trial level. For spontaneous gaze shifts, head movement onset fell along a continuum, with the majority of attempted head movements beginning before saccade onset (70%), with a median latency of 90 ms before saccade onset (Fig. 3E, G). In contrast, for stimulus-evoked gaze shifts, the vast majority of attempted head movements began after saccade onset (74%), with a median latency of 30 ms after saccade onset (Fig. 3F,G).

We next examined the amplitudes of head and eye movements during spontaneous and stimulus-evoked gaze shifts. During spontaneous gaze shifts, attempted head rotations were in the same direction as saccades and scaled with saccade amplitude (Fig. 3A, B, H). These data are consistent with eye-head coupling patterns previously observed in both head-fixed and freely moving conditions (Meyer et al., 2020). Similarly, stimulus-evoked gaze shifts involved attempted head rotations that were made in the same direction as saccades and scaled with saccade amplitude (Fig.3A, B, H). Interestingly, stimulus-evoked saccades of a given amplitude were coupled to an average of 37% smaller attempted head movements than were spontaneous saccades, due largely to the lack of slow pre-saccadic head movements observed during spontaneous gaze shifts (linear regression slopes 0.135 vs. 0.214, p < 10^-5^, permutation test). To confirm that the differences in coupling we observed were not an artifact of differences in saccade size and starting position between saccade types, we performed an additional analysis using subsets of gaze shifts matched for saccade amplitude and initial eye position and observed the same effects (Supplementary Fig. 7; linear regression slopes 0.135 vs. 0.224, p < 10^-5^, permutation test).

Because touch-evoked saccades and attempted head movements are proportional, and our previous analyses showed that saccade direction and amplitude depend on initial eye position (Fig. 2E-H), we asked whether there was a relationship between airpuff-evoked attempted head movements and initial eye position. We analyzed whisker airpuff-evoked gaze shifts because these involved a mixture of saccade directions. Strikingly, both saccade and attempted head movement direction and amplitude were dependent on initial eye position (Supplementary Fig. 8).

### The superior colliculus mediates airpuff-evoked gaze shifts

We next sought to identify the neural circuitry underlying airpuff-evoked gaze shifts. As discussed previously, in other species, stimulus-evoked gaze shifts involving directed head and eye movements are driven by SC (Freedman, 2008; Freedman et al., 1996; Guitton, 1992; Guitton et al., 1980; Paré et al., 1994). In contrast, it is widely believed that the recentering saccades observed in mice are driven by brainstem circuitry in response to head rotation (Curthoys, 2002; Hepp et al., 1993; Kitama et al., 1995; Meyer et al., 2020; Michaiel et al., 2020; Payne and Raymond, 2017). To determine whether SC is required to generate touch-evoked gaze shifts in mice, we pursued an optogenetic strategy to perturb SC activity in the period surrounding airpuff onset. For inhibition experiments, we stereotaxically injected adeno-associated virus (AAV) encoding the light-gated chloride pump eNpHR3.0 under the control of a pan-neuronal promoter and implanted a fiber optic in right SC (Gradinaru et al., 2010). Consistent with data in foveate species (Hikosaka and Wurtz, 1985; Robinson, 1972; Schiller and Stryker, 1972), optically reducing right SC activity shifted airpuff-evoked saccade endpoints to the right (i.e., ipsilaterally) for both left (−3.7 ± 4.3° (control) vs. −2.3 ± 4.7° (LED on), p < 0.001, Welch’s t-test) and right ear airpuffs (4.5 ± 4.1° (control) vs. 5.4 ± 4.7° (LED on), p = 0.011, Welch’s t-test) (Fig. 4A-C). To control for potential mismatches in starting eye position between LED-off and LED-on trials, we performed additional analyses using matched trials and found that the endpoint and amplitude differences persisted (Supplementary Fig. 10). For stimulation experiments, we stereotaxically injected AAV encoding the light-gated ion channel ChR2 under the control of a pan-neuronal promoter and implanted a fiber optic in right SC (Gradinaru et al., 2010). Because strong SC stimulation can evoke saccades, we used weak stimulation (50-120 μW) in order to bias SC activity. This manipulation caused the reciprocal effect of right SC inhibition, biasing endpoints leftwards (*i.e.,* contraversively) for both left (−5.8 ± 4.6° (control) vs. −8.0 ± 6.1° (LED on), p = 0.0016, Welch’s t-test) and right (5.6 ± 3.5° (control) vs. 1.9 ± 6.1° (LED on), p < 10^-5^, Welch’s t-test) airpuffs (Fig. 4E-G). Once again, controlling for differences in starting eye position between conditions yielded similar results (Supplementary Fig. 10).

**Figure 4.**
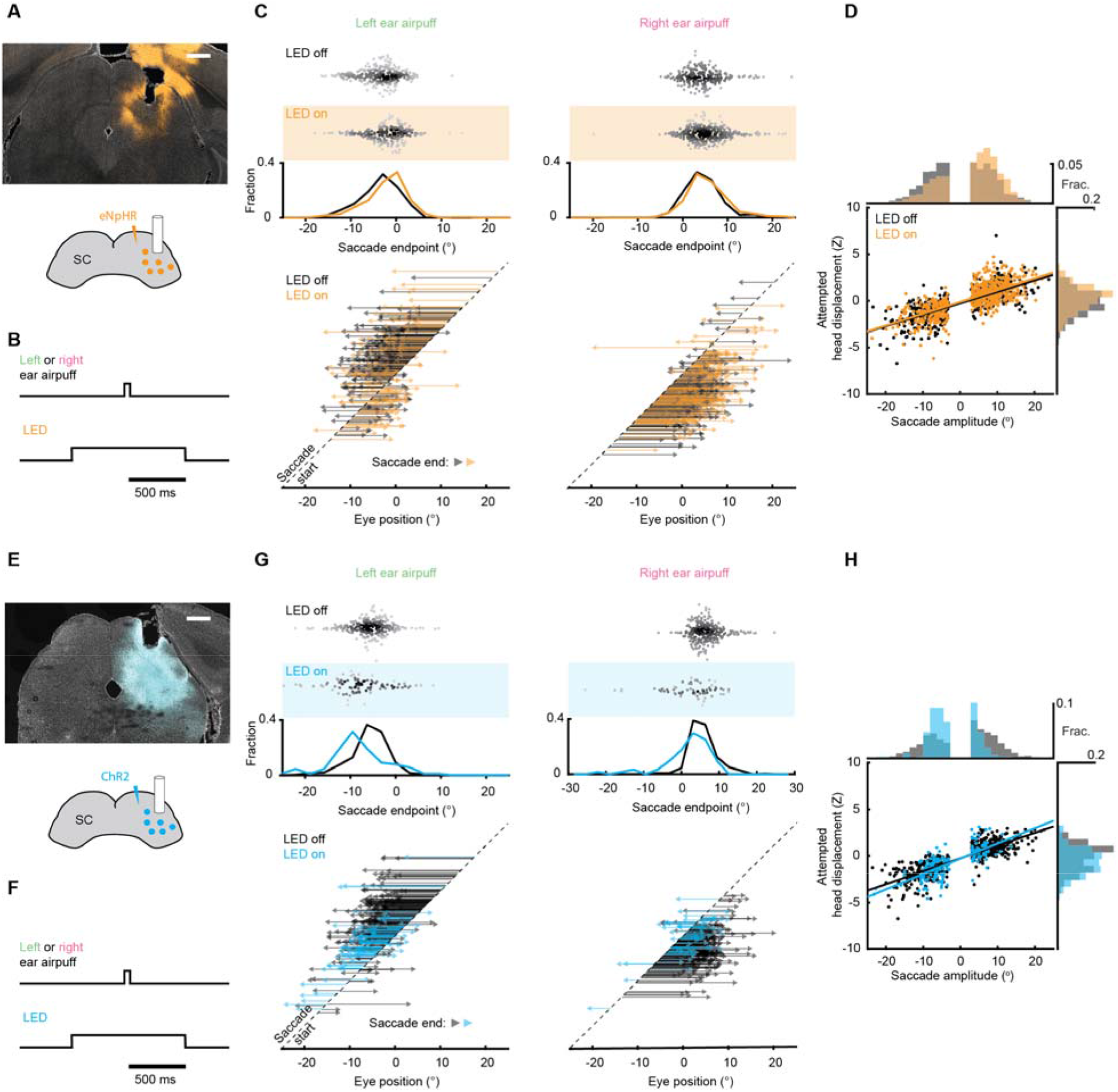
Superior colliculus controls touch-evoke gaze shifts. **(A)** Schematic of right SC optogenetic inhibition using eNpHR3.0 and example histology for representative mouse. Scale bar, 0.5 mm. The lack of fluorescence immediately surrounding fiber tip is due to photobleaching by high photostimulation intensity (12 mW, as opposed to 50-120 μW for ChR2 experiments in (E-H)). **(B)** Trial structure. Optogenetic illumination is provided for a 1 s period centered around airpuff delivery. **(C)** Effects of SC optogenetic inhibition on saccade endpoints. Top, scatter plots and histograms of endpoints for control (white background, n = 296) and LED on (orange background, n = 235) trials. Middle, endpoint histograms for control (black) and LED on (orange) trials. Bottom, saccade vectors for control (black) and LED on (orange) trials. **(D)** Head-eye amplitude coupling during ear airpuff-evoked gaze shifts for control (black) and LED on (orange) trials. Each dot represents an individual gaze shift. Control: R^2^ = 0.56, slope = 0.123, p < 10^-10^. LED on: R^2^ = 0.53, slope = 0.127, p < 10^-10^. Control and LED on regression slopes were significantly different (p=0.01, permutation test) due to differences in eye positions from which gaze shifts were generated, because controlling for initial eye position eliminated this difference (Supplementary Fig. 10). Histograms above and beside scatter plot show distributions of saccade amplitudes and attempted head displacements, respectively. Distribution means were significantly different (p < 10^-5^, permutation test). **(E)** Schematic of right SC optogenetic subthreshold stimulation using ChR2 and example histology for representative mouse. Scale bar, 0.5 mm. **(F)** Trial structure. Optogenetic illumination is provided for a 1 s period centered around airpuff delivery. **(G)** Effects of weak SC optogenetic stimulation on saccade endpoints. Top, scatter plots and histograms of endpoints for control (white background, n = 547) and LED on (blue background, n = 157) trials. We observed fewer trials in the LED-on condition because SC stimulation increased the probability of spontaneous saccades prior to stimulus onset, and trials with saccades in the 500 ms before stimulus delivery were excluded from analysis. Middle, histograms of endpoints for control (black) and LED on (blue) trials. Bottom saccade vectors for control (black) and LED on (blue) trials. **(H)** Head-eye amplitude coupling during ear airpuff-evoked gaze shifts for control (black) and LED on (blue) trials. Each dot represents an individual gaze shift. Attempted head amplitude was measured 150 ms after saccade onset. Control: R^2^ = 0.69, slope = 0.137, p < 10^-10^. LED on: R^2^ = 0.52, slope = 0.164, p < 10^-10^. Control and LED-on regression slopes were significantly different (p < 10^-5^, permutation test) due to difference in eye positions from which gaze shifts were generated, because controlling for initial eye position eliminated this difference (Supplementary Fig. 10). Histograms above and beside scatter plot show distributions of saccade amplitudes and attempted head displacements, respectively. Distribution means were significantly different (p < 10^-5^, permutation test).

To understand how SC manipulations affect attempted head movements and head-eye coupling, we examined the distribution of head movements as a function of saccade amplitude. As expected given the role of SC in generating both head and eye movements, attempted head movements were shifted to the right (*i.e.,* ipsiversively) by SC inhibition (Fig. 4D) and to the left (*i.e.,* contraversively) by SC excitation (Fig. 4H). To examine the effects of SC manipulations on head-eye coupling, we identified trials with identical saccade trajectories in LED-on and LED-off conditions and examined the corresponding head movement amplitudes. Interestingly, SC manipulations had no effect on the relationship between saccade and head movement amplitudes, suggesting that SC manipulations do not change head-eye coupling during gaze shifts. Taken together, these bidirectional manipulations indicate that SC serves a conserved necessary and sufficient role in generating ear airpuff-evoked gaze shifts in which it specifies overall gaze shift amplitude.

## Discussion

Here we investigated whether mouse gaze shifts are more diverse than had previously been appreciated. In the prevailing view, mouse gaze shifts are led by head rotations that trigger compensatory eye movements, including saccades that function to reset the eyes (Land, 2019; Land and Nilsson, 2012; Liversedge et al., 2011; Meyer et al., 2020; Michaiel et al., 2020; Payne and Raymond, 2017). These “recentering” saccades are attributed to head movement-related vestibular and optokinetic cues (Curthoys, 2002; Meyer et al., 2020; Michaiel et al., 2020; Payne and Raymond, 2017). Working in a head-fixed context to eliminate vestibular and optokinetic cues and to present stimuli of different modalities at precise craniotopic locations, we found that mice are capable of an additional type of gaze shift. As discussed below, we identified numerous features that distinguish this new type of gaze shift from the previously studied type, including the endpoints of the saccades, their timing and amplitude relative to attempted head movements, and the brain regions that drive them.

### Touch-evoked saccades are directed

The first indication that touch-evoked gaze shifts differ from those previously observed in mice was an endpoint analysis revealing that touch-evoked saccades are directed rather than recentering. This conclusion is based on three lines of evidence. First, endpoints of saccades evoked by left and right ear airpuffs are near the left and right edges, respectively, of the range of eye positions observed and overlap minimally, despite trial-to-trial variability. In contrast, endpoints for saccades evoked by left and right auditory stimuli are indistinguishable and well described by the recentering model. Second, left and right ear airpuffs evoke saccades traveling in opposite directions from most eye positions; by definition, one of these directions must lead away from center and is thus *centrifugal* rather than *centripetal*. In contrast, saccades evoked by both left and right auditory stimuli travel centripetally from all initial eye positions. Third, from many eye positions, touch-evoked saccades that begin towards the center pass through to reach endpoints at eccentricities between 5 to 10 degrees and cannot accurately be termed centripetal. For these reasons, we conclude that touch-evoked saccades are directed and do not serve to recenter the eyes.

Our findings contrast with and complement previous studies contending that rodents, like other afoveates, use saccades to reset their eyes to more central locations (Meyer et al., 2018, 2020; Michaiel et al., 2020; Wallace et al., 2013). One recent analysis suggested that gaze shifts made during visually guided prey capture involve resetting centripetal saccades that “catch up” with the head (Michaiel et al., 2020). Another found that saccades away from the nose recenter the eye, whereas saccades toward the nose move the eye slightly beyond center (Meyer et al., 2020). Although we observed this as well, it does not contribute to our results because we averaged the positions of the left and right eyes, eliminating this asymmetry. Earlier studies in head-fixed mice observed occasional, undirected saccades in response to changes in the visual environment (Samonds et al., 2018) and found that mice could be trained to produce visually guided saccades only after weeks of training and at extremely long (∼1 s) latencies (Itokazu et al., 2018). To our knowledge, ours is the first study demonstrating innate gaze shifts involving directed saccades in mice (or any species lacking a fovea).

### Touch-evoked gaze shifts are not led by head movements

The prevailing view holds that head movements initiate and determine the amplitude of mouse gaze shifts, with eye movements compensatory by-products. In support of this model, one study found that spontaneous saccades in head-fixed mice are preceded by attempted head rotations. A careful comparison with gaze shifts occurring during visually guided object tracking and social interactions in freely moving mice led the authors to suggest that head-eye coupling is not disrupted during head-fixation, and that gaze shifts in both contexts are head-initiated (Meyer et al., 2020). Another study tracked the eyes and head during visually guided cricket hunting and found that gaze shifts are driven by the head, with the eyes following to stabilize and recenter gaze (Michaiel et al., 2020). Together, these findings have bolstered the prevailing view that afoveates such as mice generate gaze shifts driven by head movements, with eye movements compensatory by-products (Land, 2019; Land and Nilsson, 2012; Liversedge et al., 2011). However, while our findings for spontaneous saccades are consistent with those in the literature, we have shown that touch-evoked gaze shifts are initiated by saccades, a finding that contrasts with and complements earlier studies.

This finding that touch-evoked gaze shifts are initiated by saccades suggests that mouse saccades are not always a simple by-product of head movements. Additional support for this idea came from an analysis of head and eye movements as a function of eye position. If gaze shifts were determined solely by the location of the stimulus relative to the head and saccades were a compensatory by-product of this calculation, eye position should have no effect on head movement amplitude. However, we found that the amplitudes and directions of both saccades and attempted head movements evoked by saccades vary with initial eye position. The influence of eye position on touch-evoked eye *and* head movements further indicates that saccades are not compensatory by-products of head movements. Instead, touch-evoked head movements and directed saccades are specified simultaneously as parts of a coordinated movement whose component movements take into account both stimulus location and initial eye position.

The relative timing of head and eye components during touch-evoked gaze shifts in mice resembles that observed during gaze shifts in cats and primates. For example, head-fixed cats and primates generate gaze shifts using directed saccades and then maintain their eyes in the new orbital position, similar to what we have observed in mice (Freedman, 2008; Guitton et al., 1980). In addition, saccades in head-fixed cats and primates are often accompanied by attempted head rotations, similar to those we observe during touch-evoked gaze shifts in head-fixed mice (Bizzi et al., 1971; Guitton et al., 1984; Paré et al., 1994). In primates and cats able to move their heads, gaze shifts are usually led by directed saccades (with some exceptions), likely because the eyes have lower rotational inertia and can move faster (Pelisson and Guillaume, 2009; Ruhland et al., 2013). These saccades tend to be followed by a head movement in the same direction that creates vestibular signals that drive slow, centripetal counterrotation of the eyes to maintain fixation (Bizzi et al., 1972; Freedman, 2008; Freedman and Sparks, 1997; Guitton et al., 1984). In this way, the animal can rapidly shift its gaze with a directed saccade yet subsequently reset the eyes to a more central position as the head moves. It is tempting to speculate that a similar coordinated sequence of head and eye movements occurs during freely moving touch-evoked gaze shifts, enabling mice to rapidly shift gaze with their eyes while eventually resetting the eyes in a more central orbital position.

### Head-eye amplitude coupling differs during spontaneous and evoked saccades

An additional feature that distinguishes spontaneous and touch-evoked gaze shifts is the relative contributions of head and eye movements. We found that spontaneous saccades of a given amplitude are coupled to larger head movements than are touch-evoked saccades. This difference arises largely from the absence of a pre-saccadic attempted head movement during touch-evoked gaze shifts. This differential pairing of head and eye movements is reminiscent of reports in primates and cats that the relative contributions of head and eye movements vary for gaze shifts evoked by different sensory modalities (Goldring et al., 1996; Populin, 2006; Populin and Rajala, 2011; Populin et al., 2004a; Ruhland et al., 2013; Tollin et al., 2005). However, in those species, vision typically elicits gaze shifts dominated by saccades while hearing typically evokes gaze shifts entailing larger contributions from head movements. In contrast, we observed that sound- and touch-evoked gaze shifts involve larger contributions from saccades than do spontaneous gaze shifts, whereas visual stimuli did not evoke gaze shifts at all. This indicates that although there is general conservation of the involvement of SC in sensory-driven gaze shifts, modality-specific features are not conserved, which may reflect differences in sensory processing across species. For example, virtually every cell in primate intermediate and deep SC that is responsive to tactile stimuli is also responsive to visual stimuli (Groh and Sparks, 1996). In contrast, a recent study of *Pitx2*^+^ neurons in the intermediate and deep mouse SC, which project to brainstem oculomotor centers and when optogenetically stimulated evoke orienting movements of the eyes and head reminiscent of the gaze shifts we describe (Masullo et al., 2019), reported that these neurons responded robustly to whisker airpuffs but did not respond to visual stimuli (Xie et al., 2021). We also observed limited variability across mice in the relative contributions of head and eye movements to gaze shifts, which contrasts with the observation that different human subjects are head “movers” and “non-movers” during gaze shifts (Supplementary Fig. 9) Thus, our results reveal that mice are capable of using multiple strategies to shift their gaze but with key differences from other species.

### The role of superior colliculus in sensory-evoked mouse gaze shifts

In other species, SC drives sensory-evoked gaze shifts, and microstimulation and optogenetic stimulation of mouse SC has been shown to elicit gaze shifts (Masullo et al., 2019; Wang et al., 2015). However, to our knowledge, no study had identified a causal involvement of SC in mouse gaze shifts. We performed bidirectional optogenetic manipulations that revealed that touch-evoked gaze shifts depend on SC, identifying a conserved, necessary and sufficient role for SC in directed gaze shifts. In addition, we found that SC manipulations did not alter head-eye amplitude coupling. This observation suggests that SC specifies the overall gaze shift amplitude rather than the individual eye or head movement components, consistent with observations in other species (Freedman et al., 1996; Paré et al., 1994).

### Ethological significance

Prior to the present study, it was believed that species with high-acuity retinal specializations acquired the ability to make directed saccades to scrutinize salient environmental stimuli, because animals lacking such retinal specializations were thought incapable of gaze shifts led by directed saccades (Land, 2019; Land and Nilsson, 2012; Liversedge et al., 2011; Walls, 1962). Our discovery that sensory-guided directed saccades are present in mice—albeit without the precise targeting of stimulus location seen in foveate species—raises the question of what fovea-independent functions these movements serve. Although mice have lateral eyes and a large field of view, saccades that direct gaze towards a stimulus, as seems to occur with both ear and whisker tactile stimuli, may facilitate keeping salient stimuli within the field of view. As natural stimuli are often multimodal, directing non-visual stimuli towards the center of view maximizes the likelihood of detecting the visual component of the stimulus. Alternatively, despite mouse retinae lacking discrete, anatomically defined specializations such as foveae or areas centralis, there are subtler nonuniformities in the distribution and density of photoreceptors and retinal ganglion cell subtypes, and magnification factor, receptive field sizes, and response tuning vary across the visual field in higher visual centers; it may be desirable to center a salient tactile stimulus on a particular retinal region to enable scrutiny of the visual component of the stimulus (Ahmadlou and Heimel, 2015; Baden et al., 2013; van Beest et al., 2021; Bleckert et al., 2014; Drager and Hubel, 1976; Feinberg and Meister, 2015; Li et al., 2020; de Malmazet et al., 2018). Although touch-evoked saccades alone may be too small to center the stimulus location on any particular region of the retina, they may do so in concert with directed head movements.

Why tactile stimuli evoked directed saccades in our preparation whereas auditory and visual stimuli do not is unclear. One possibility is the aforementioned speed of saccades relative to head movements may be especially beneficial because tactile stimuli typically derive from proximal objects and as a result may demand rapid responses. Alternatively, because the spatial acuity of the tactile system is higher than that of the auditory system in mice, this may enable more precise localization of the tactile stimuli (Allen and Ison, 2010; Diamond et al., 2008). Auditory stimuli, in contrast, may alert animals to the presence of a salient stimulus in their environments whose location is less precisely ascertained, and as a result drive gaze shifts whose goal is to reset the eyes to a central position that maximizes their chances of sensing and responding appropriately. Finally, our set of stimuli was not exhaustive, and it is possible that as yet unidentified visual or auditory stimuli could elicit gaze shifts with directed saccades. Alternatively, the recent report that mouse SC *Pitx2^+^* neurons respond to tactile but not visual stimuli may indicate that visual orienting is driven by a distinct neural pathway or that head-fixing gates visual but not tactile responses of SC neurons.

### Future Directions

In this study we used a head-fixed preparation to eliminate the confound of head movement-related sensory cues and to present stimuli from defined locations. However, in the future, it will be interesting to compare touch-evoked gaze shifts in head-fixed and freely moving animals. For example, as noted previously, whereas head-fixed primates and cats generate gaze shifts using directed saccades and then maintain their eyes in the new orbital position, similar to what we have observed, in freely moving primates and cats, gaze shifts are led by directed saccades but typically followed by head movements during which the eyes counterrotate centripetally in order to maintain gaze in the new direction. It will be interesting to know whether similar differences distinguish saccade-led touch-evoked gaze shifts in head-fixed and freely moving mice. By expanding on methods recently described by other groups, it may be possible to investigate these and other questions (Meyer et al., 2018, 2020; Michaiel et al., 2020; Payne and Raymond, 2017).

Furthermore, a practical implication of our identification of mouse SC-dependent gaze shifts is that this behavioral paradigm could be applied to the study of several outstanding questions. First, there are many unresolved problems regarding the circuitry and ensemble dynamics underlying target selection (Basso and May, 2017) and saccade generation (Gandhi and Katnani, 2011), and the mouse provides a genetically tractable platform with which to investigate these and other topics. Second, gaze shifts are aberrant in a host of conditions, such as Parkinson’s and autism spectrum disorder (Liversedge et al., 2011). This paradigm could be a powerful tool for the study of mouse models of a variety of neuropsychiatric conditions. Third, directing saccades towards particular orbital positions during these gaze shifts requires an ability to account for the initial positions of the eyes relative to the target, a phenomenon also known as remapping from sensory to motor reference frames. Neural correlates of this process have been observed in primates (Groh and Sparks, 1996; Jay and Sparks, 1984) and cats (Populin et al., 2004b), but the underlying circuitry and computations remain obscure. This behavior may facilitate future studies of this problem. Fourth, the different types of gaze shifts that rely on distinct head-eye coupling we have identified may be useful for understanding mechanisms that control movement coordination. Thus, touch-evoked saccade behavior is likely to be a powerful tool for myriad lines of investigation.

## Conclusions

We have found that mice make an unexpectedly broad range of gaze shifts, including a new type that is elicited by sensory stimuli, led by directed saccades, and incorporates smaller head movements. Prior studies in species whose retinae lack high-acuity specializations had never observed this type of gaze shift, but our study used a broader range of stimuli than previously tested in a preparation that allowed spatially precise delivery. Detailed perturbation experiments determined that the circuit mechanisms of sensory-evoked gaze shifts are conserved from mice to primates, suggesting that this behavior may have arisen in a common, afoveate ancestral species long ago. More broadly, our findings suggest that analyzing eye movements of other afoveate species thought not to make directed saccades—such as rabbits, toads, and goldfish—in response to a diverse range of multimodal stimuli may uncover similar abilities to make diverse types of gaze shifts.

## Acknowledgments

We thank M. Brainard, J. Horton, A. Krishnaswamy, M. Scanziani, and members of the Feinberg laboratory for helpful discussions and comments on earlier versions of the manuscript. This work was supported by departmental funds and grants from the E. M. Ziegler Foundation for the Blind, Sandler Foundation, Klingenstein-Simons Fellowship Award in Neuroscience, Brain and Behavior Research Foundation (NARSAD Young Investigator Awards 25337 and 27320), Whitehall Foundation, Simons Foundation (SFARI 574347), and US National Institutes of Health (DP2 MH119426 and R01 NS109060) to E.H.F.

## Author contributions

S.H.Z., D.E.T., and E.H.F. conceived of the project, designed the experiments, analyzed the data, and prepared the manuscript. S.H.Z., D.E.T., J.Y.W., and J.M.A. conducted the experiments.

---

**Supplementary Figure 1.**
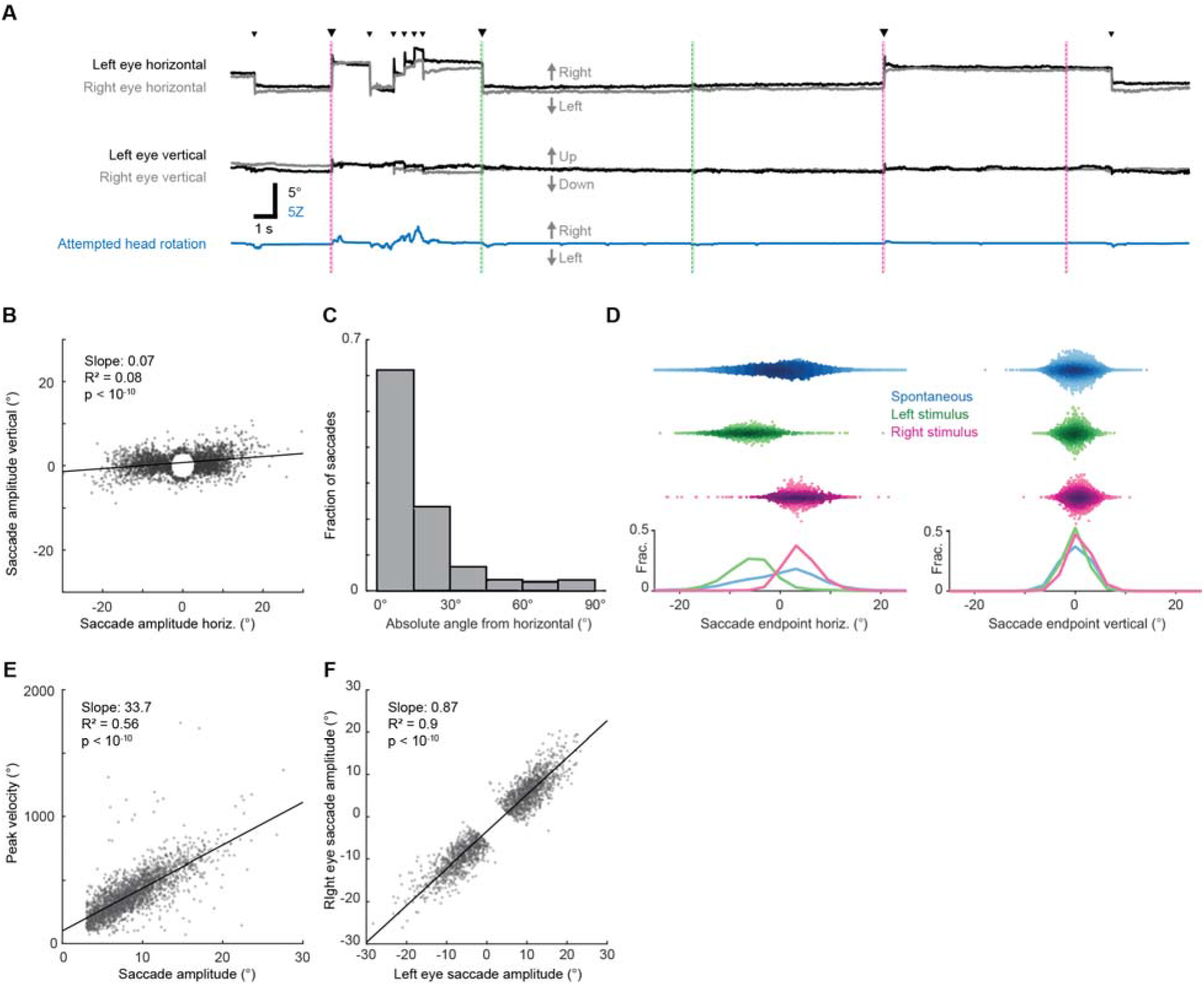
Airpuffs evoke horizontal saccades. **(A)** Sample traces showing pupil azimuth (top) and elevation (middle) and attempted head rotation (bottom). Black corresponds to left eye, gray corresponds to right eye. Small arrowheads indicate spontaneous saccades. Large arrowheads indicate stimulus-evoked directed saccades. Green and magenta dashed vertical lines correspond to left and right ear airpuffs, respectively. **(B)** Ear airpuff-evoked saccade endpoints and linear fit. n = 5 mice, 2337 trials. **(C)** Distribution of angles between airpuff-evoked saccade vectors and horizontal axis. Gray bars indicate population means. n = 5 mice, 2337 trials **(D)** Distributions of saccade endpoints in horizontal and vertical axes. n = 5 mice; trials = 16291 (spontaneous), 1067 (left airpuff), 1270 (right airpuff**). (E)** Relationship between saccade amplitude and peak velocity**. (F)** Relationship between right and left eye saccade amplitudes. n = 4 mice, 1861 trials.

**Supplementary Figure 2.**
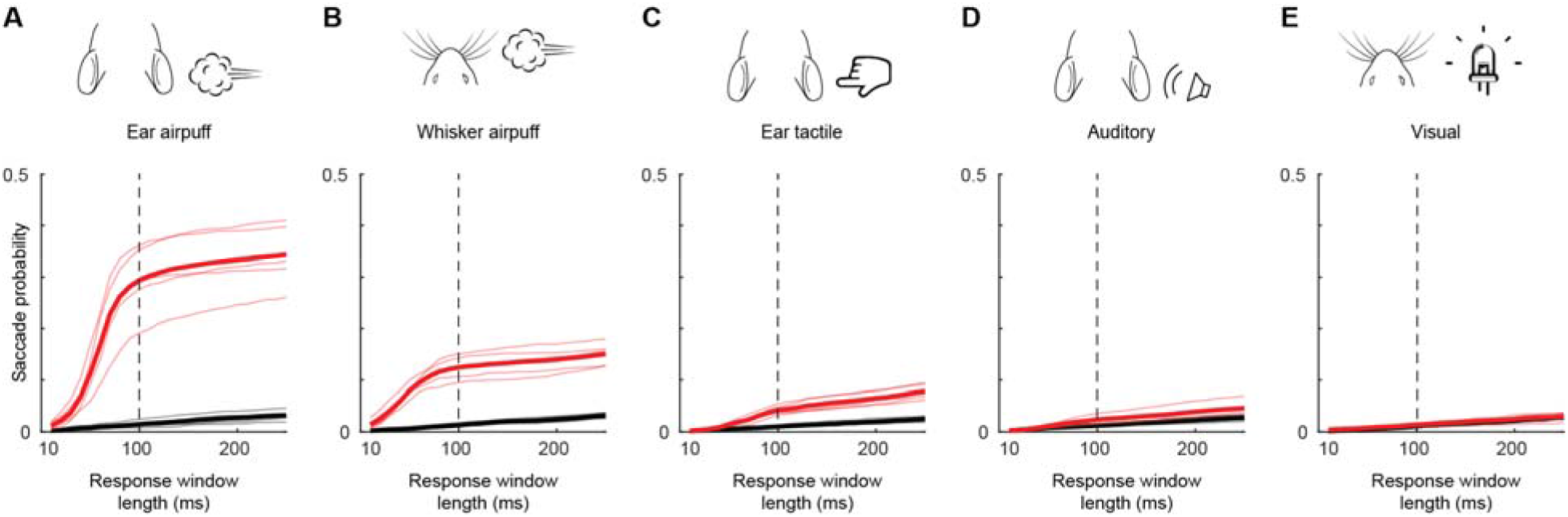
Evoked saccades occur within a narrow window after stimulus delivery. **(A-E)** Cumulative probabilities of detecting evoked (red) and spontaneous (black) saccades as a function of response window length for ear airpuffs (A), whisker airpuffs (B), ear tactile stimuli (C), auditory airpuffs (D), and visual stimuli (E). Thin lines denote values for individual mice, thick lines denote population mean. Dashed vertical line indicates end of window used for analyses of evoked saccades.

**Supplementary Figure 3.**
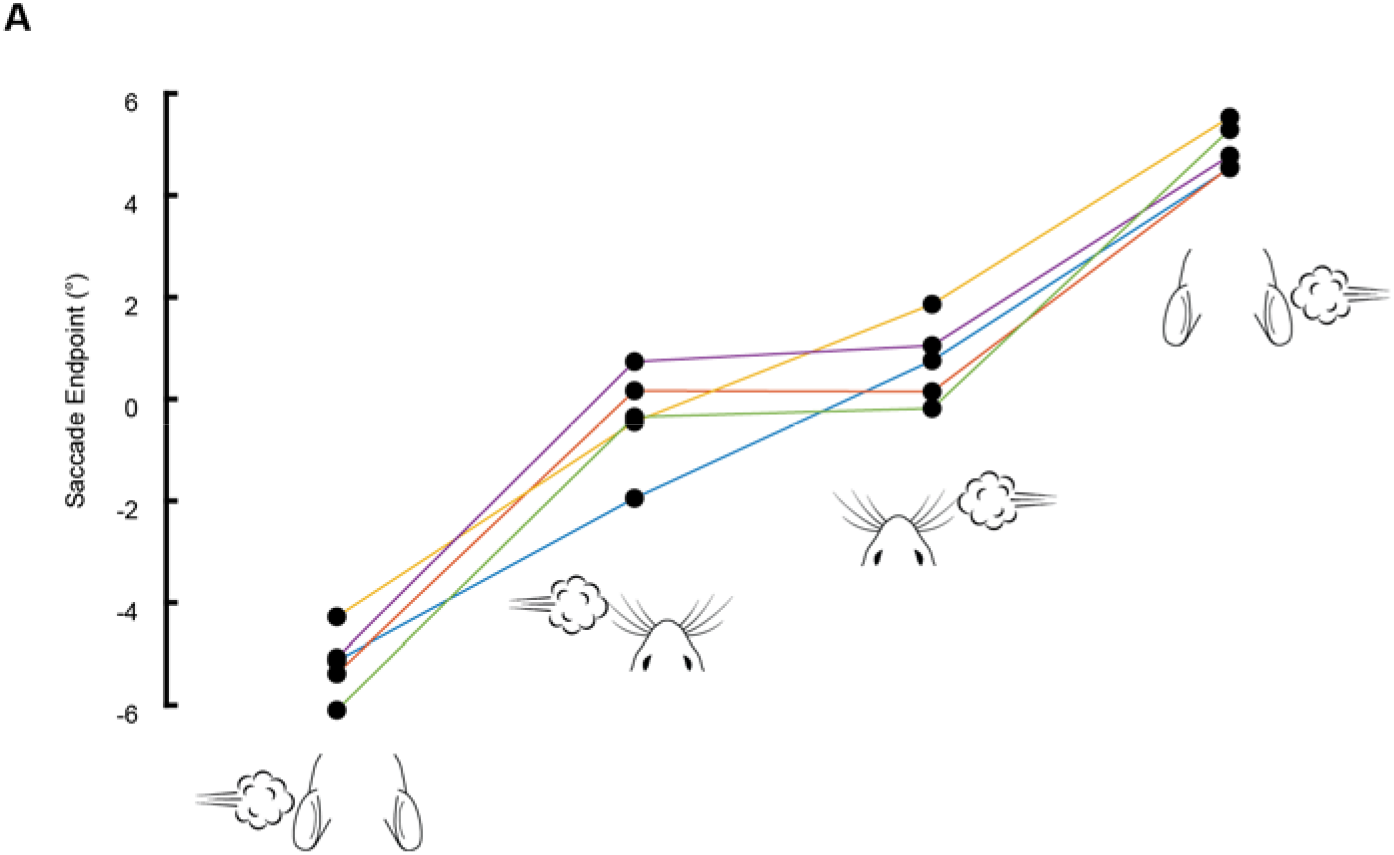
Endpoints of airpuff-evoked saccades are ordered according to site of stimulation. Each line corresponds to a single mouse and shows mean endpoint of saccades evoked by (in order, from left to right) left ear airpuffs, left whisker airpuffs, right whisker airpuffs, and right ear airpuffs).

**Supplementary Figure 4.**
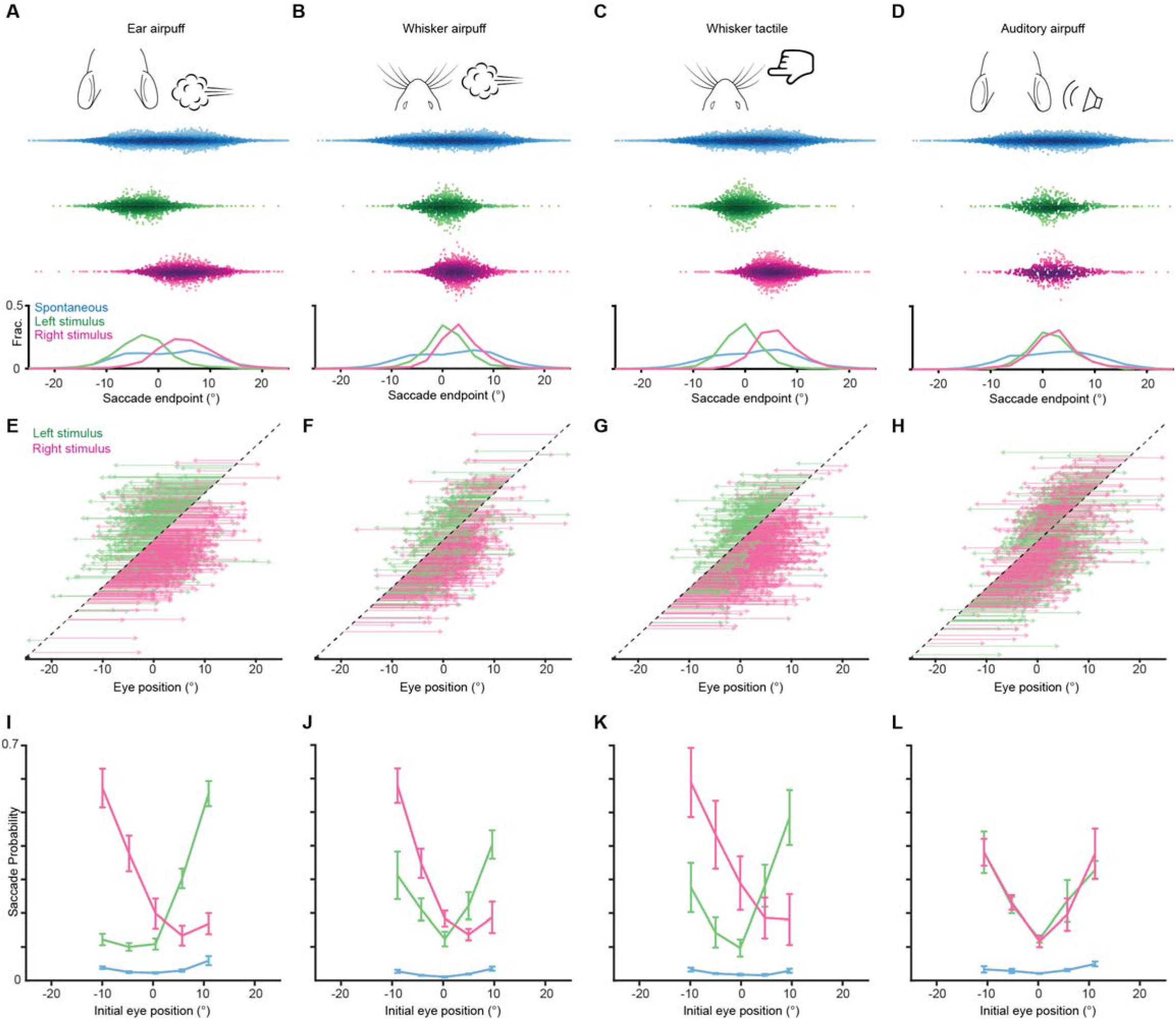
Endpoints and trajectories of sensory-evoked saccades for an additional cohort of mice. **(A-D)** Endpoints for ear airpuff-, whisker airpuff-, ear tactile-, and auditory airpuff-evoked saccades. Top, schematics of stimuli. Middle, scatter plots showing endpoints of all saccades for all animals made spontaneously (blue) and in response to left (green) and right (magenta) stimuli. Darker shading indicates areas of higher density. Bottom, endpoint distributions for spontaneous and evoked saccades. **(E-H)** Trajectories of individual stimulus-evoked saccades. Each arrow denotes the trajectory of a single saccade. Saccades are sorted according to initial eye positions, which fall on the dashed diagonal line. Saccade endpoints are indicated by arrowheads. Because the probability of evoked gaze shifts differed across stimuli, data are randomly subsampled to show roughly equal numbers of trials for each condition. **(I-L)** Relationship between eye position and saccade probability. Green and magenta lines indicate population means for saccades evoked by left and right stimuli, respectively. Blue lines indicate spontaneous saccades. Error bars indicate s.e.m Saccade numbers for A-L: ear airpuff sessions, spontaneous = 14304, left ear airpuff-evoked = 1221 (244 in E), right ear airpuff-evoked = 1755 (351 in E); whisker airpuff sessions, spontaneous = 8971, left whisker airpuff-evoked = 1107 (221 in F), right whisker airpuff-evoked = 1482 (296 in F); whisker tactile sessions, spontaneous = 13242, left whisker-evoked = 1473 (294 in G), right whisker-evoked = 2408 (481 in G); auditory sessions, spontaneous = 8774, left auditory-evoked = 833 (333 in H), right auditory-evoked = 757 (302 in H).

**Supplementary Figure 5.**
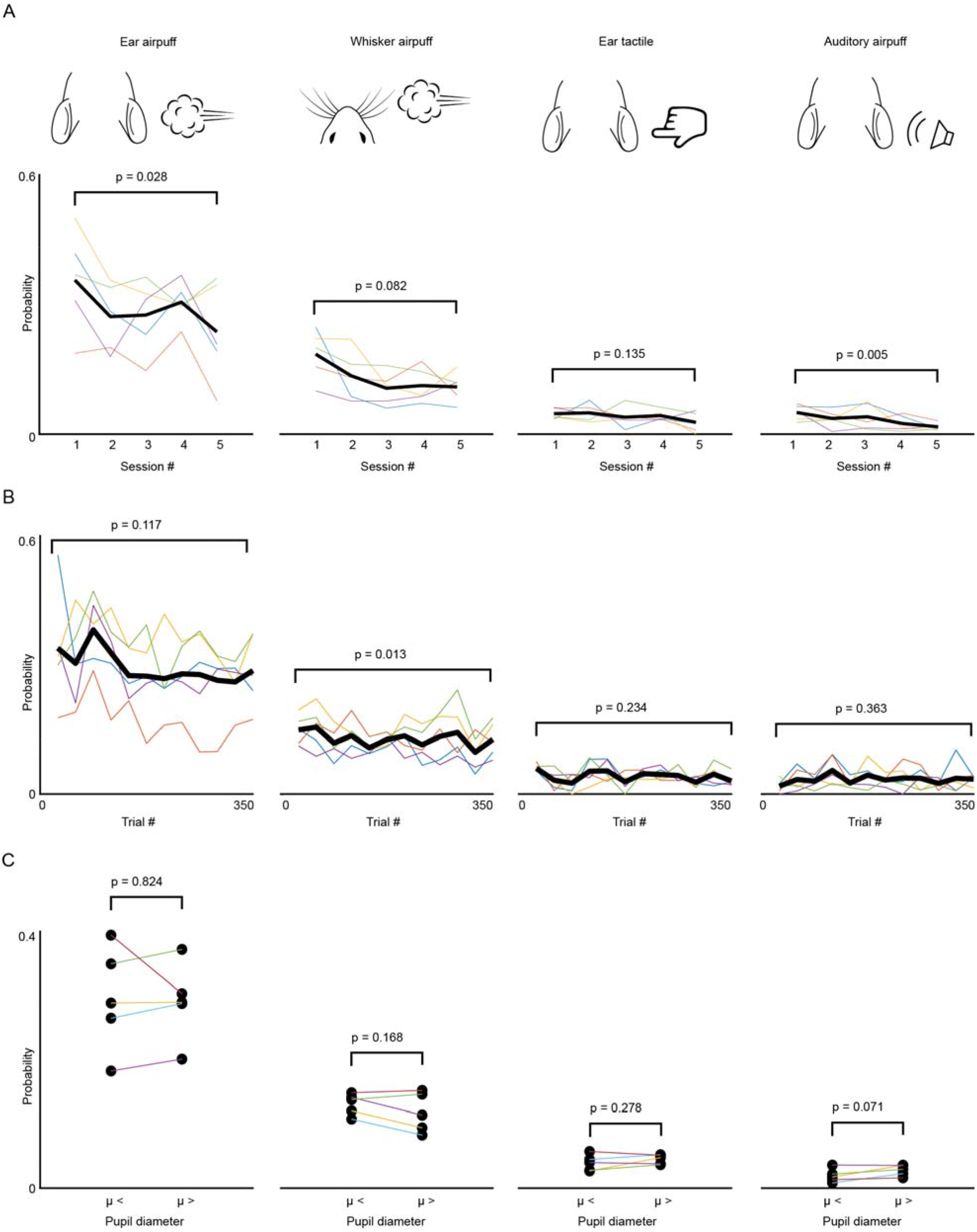
Effects of sensory history and arousal on saccade generation. (**A**) Overall gaze shift probability across 5 sessions for ear airpuffs, whisker airpuffs, ear tactile, and auditory airpuffs stimuli. Each thin colored line corresponds to an individual mouse. Black line corresponds to mean (**B**) As in (A) but for gaze shift probability within sessions. **(C)** Effects of arousal on saccade probability. μ denotes mean pupil diameter. Statistical significance assessed using paired Student’s t-test.

**Supplementary Figure 6.**
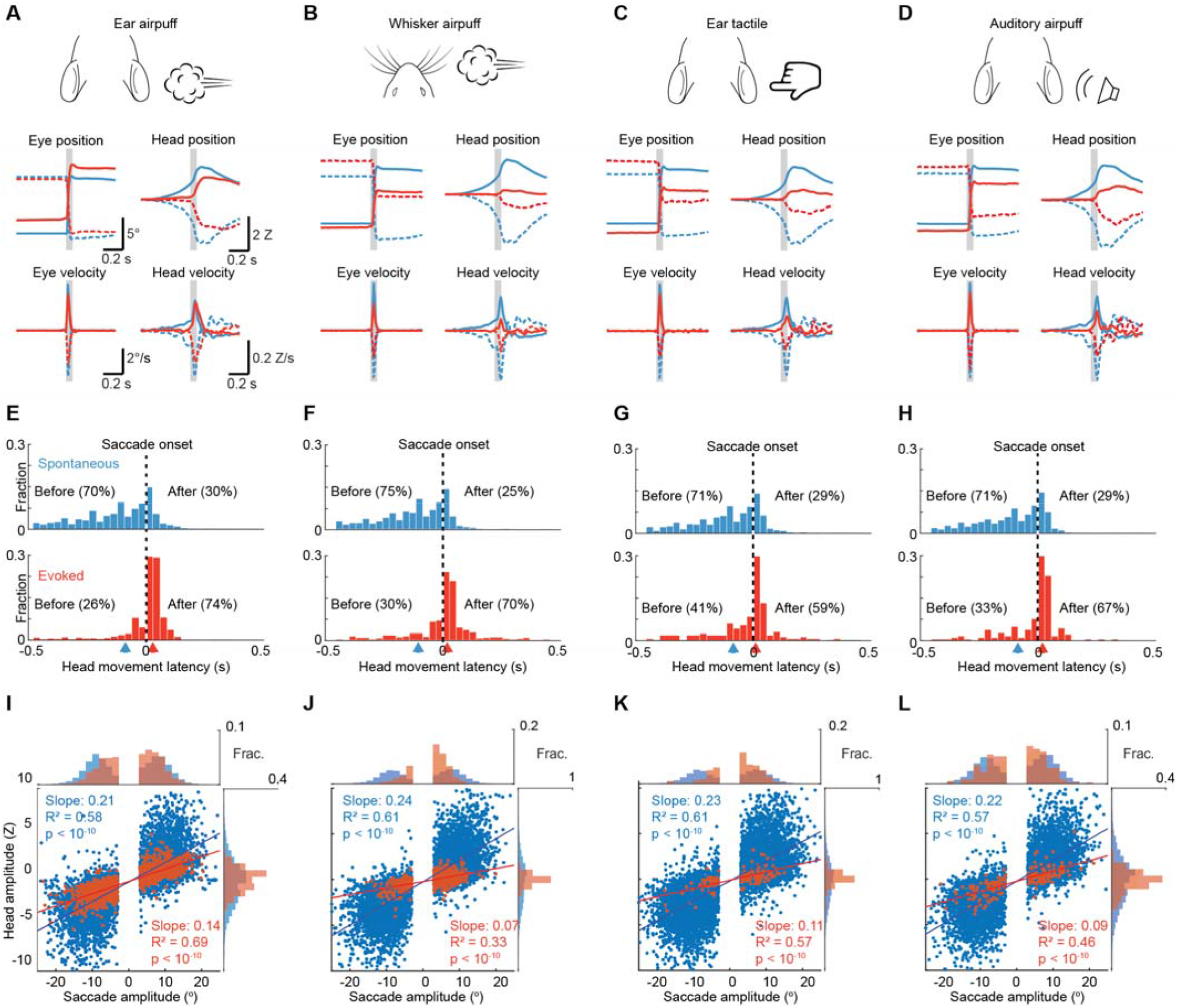
Head-eye coupling for different stimuli. **(A)** Top, stimulus schematics. Middle left, mean trajectories of rightward (solid traces) and leftward (dashed traces) saccades during spontaneous (blue, n= 7146) and ear airpuff-evoked (red, n = 1437) gaze shifts. Middle right, mean attempted head displacement accompanying rightward (solid traces) and leftward (dashed traces) saccades during spontaneous (blue) and ear airpuff-evoked (red) gaze shifts. Bottom left, mean velocities of all rightward (solid traces) and leftward (dashed traces) saccades during spontaneous (blue) and ear airpuff-evoked (red) gaze shifts. Bottom right, mean attempted head movement velocities accompanying rightward (solid traces) and leftward (dashed traces) saccades during spontaneous (blue) and ear airpuff-evoked (red) gaze shifts. **(B-D)** As in (A) for whisker airpuffs (B), ear tactile (C) and auditory airpuffs (D). n = 5 mice. Trial numbers: whisker airpuff sessions (spontaneous = 7790, evoked = 628), ear tactile sessions (spontaneous = 6706, evoked = 187), auditory airpuff sessions (spontaneous = 10240, evoked = 148). (**E-H)** Distributions of attempted head movement latencies relative to saccade onset for spontaneous (top) and evoked (bottom) saccades. Medians indicated by blue and red arrowheads (ear airpuff sessions, spontaneous: −90 ms, evoked: 30 ms; whisker airpuff sessions, spontaneous: −110 ms, evoked: 20 ms; ear tactile sessions, spontaneous: - 90 ms, evoked: 20 ms; auditory sessions, spontaneous: −90ms, evoked: 10 ms) For each condition, distributions are significantly different for spontaneous and evoked gaze shifts (p < 10^-5^, permutation test) **(I-L)**. Head-eye amplitude coupling of spontaneous (blue) and evoked saccades (red). Each dot corresponds to a single gaze shift. Regression statistics in figure. For every stimulus type, spontaneous and evoked regression slopes were significantly different (p < 10^-5^, permutation test). Histograms above and beside scatter plot indicate distributions of saccade and attempted head movement amplitudes, respectively. For each condition, distributions were significantly different between spontaneous and evoked (p < 10^-5^, permutation test).

**Supplementary Figure 7.**
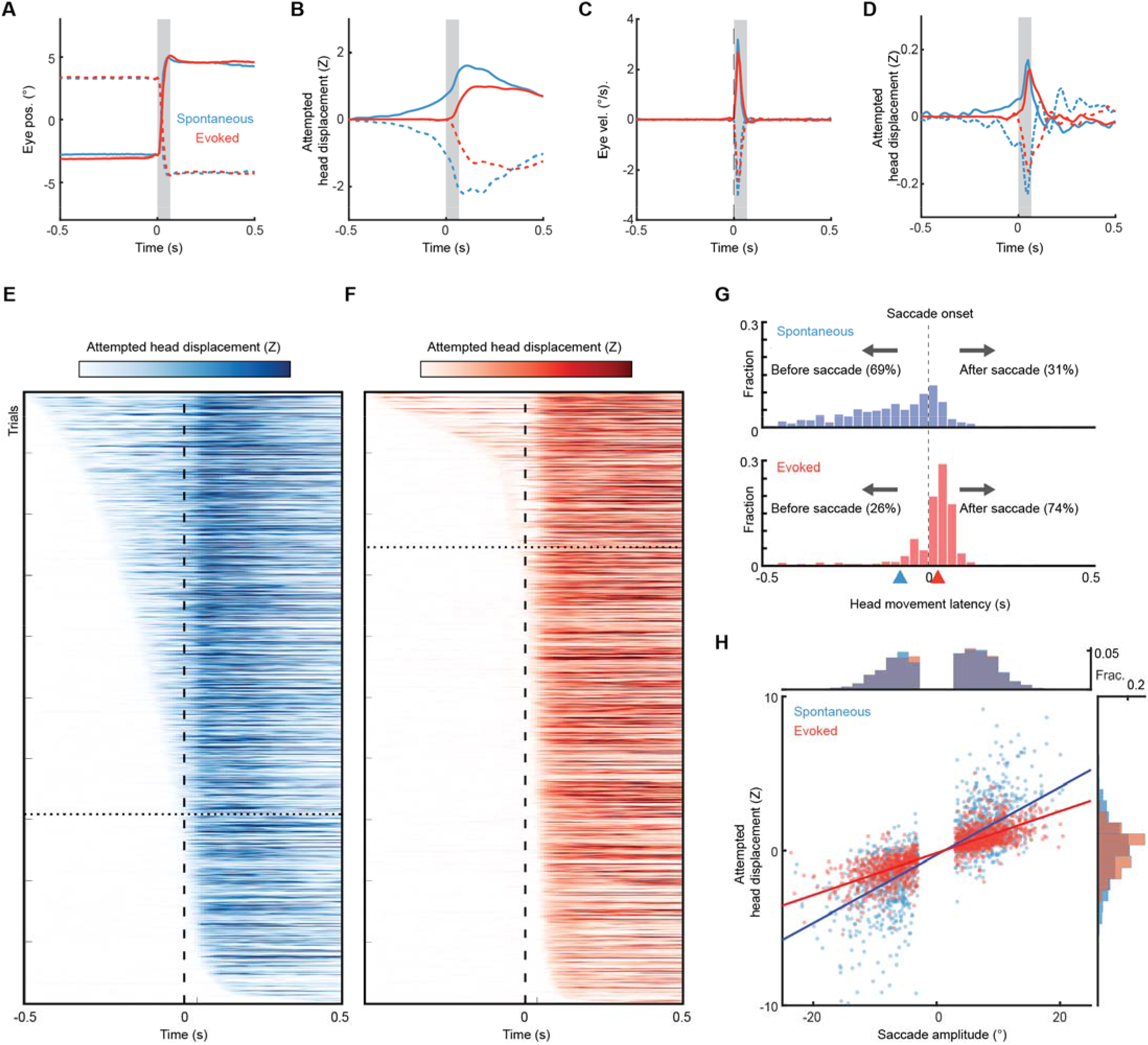
Different head-eye coupling during spontaneous and touch-evoked gaze shifts is not due to differences in gaze shift amplitudes. **(A)** Mean trajectories of amplitude and initial position-matched rightward (solid traces) and leftward (dashed traces) saccades during spontaneous (blue, n= 1437) and ear airpuff-evoked (red, n = 1437) gaze shifts. **(B)** Mean attempted head movement amplitudes accompanying rightward (solid traces) and leftward (dashed traces) saccades during spontaneous (blue) and ear airpuff-evoked (red) gaze shifts. **(C)** Mean velocities of all rightward (solid traces) and leftward (dashed traces) saccades during spontaneous (blue) and ear airpuff-evoked (red) gaze shifts. (**D**) Mean head movement velocities accompanying rightward (solid traces) and leftward (dashed traces) saccades during spontaneous (blue) and ear airpuff-evoked (red) gaze shifts. **(E, F)** Timing of attempted head movements relative to saccades during all spontaneous (E) and ear airpuff-evoked (F) gaze shifts. Each row corresponds to a single gaze shift. Darker shades indicate larger attempted head displacement. Dashed vertical line indicates time of saccade onset. Trials are sorted by latency of attempted head movements. Trials above and below dashed horizontal line correspond to attempted head movements that began before and after the saccade, respectively. **(G)** Distributions of attempted head movement latencies relative to saccade onset for spontaneous (top) and ear airpuff-evoked (bottom) saccades. Arrowheads indicate median latencies. **(H)**. Head-eye amplitude coupling of spontaneous (blue) and ear airpuff-evoked saccades (red). Each dot corresponds to a single gaze shift. Attempted head amplitude was measured 150 ms after saccade onset. Spontaneous: R^2^ = 0.60, slope = 0.224, p < 10^-10^. Evoked: R^2^ = 0.69, slope = 0.135, p < 10^-10^. Spontaneous and evoked regression slopes were significantly different (p < 10^-5^, permutation test). Histograms above and beside scatter plot indicate distributions of saccade and attempted head movement amplitudes. Difference in means were not significant (p = 0.46 for saccades, p = 0.31 for head, permutation test).

**Supplementary Figure 8.**
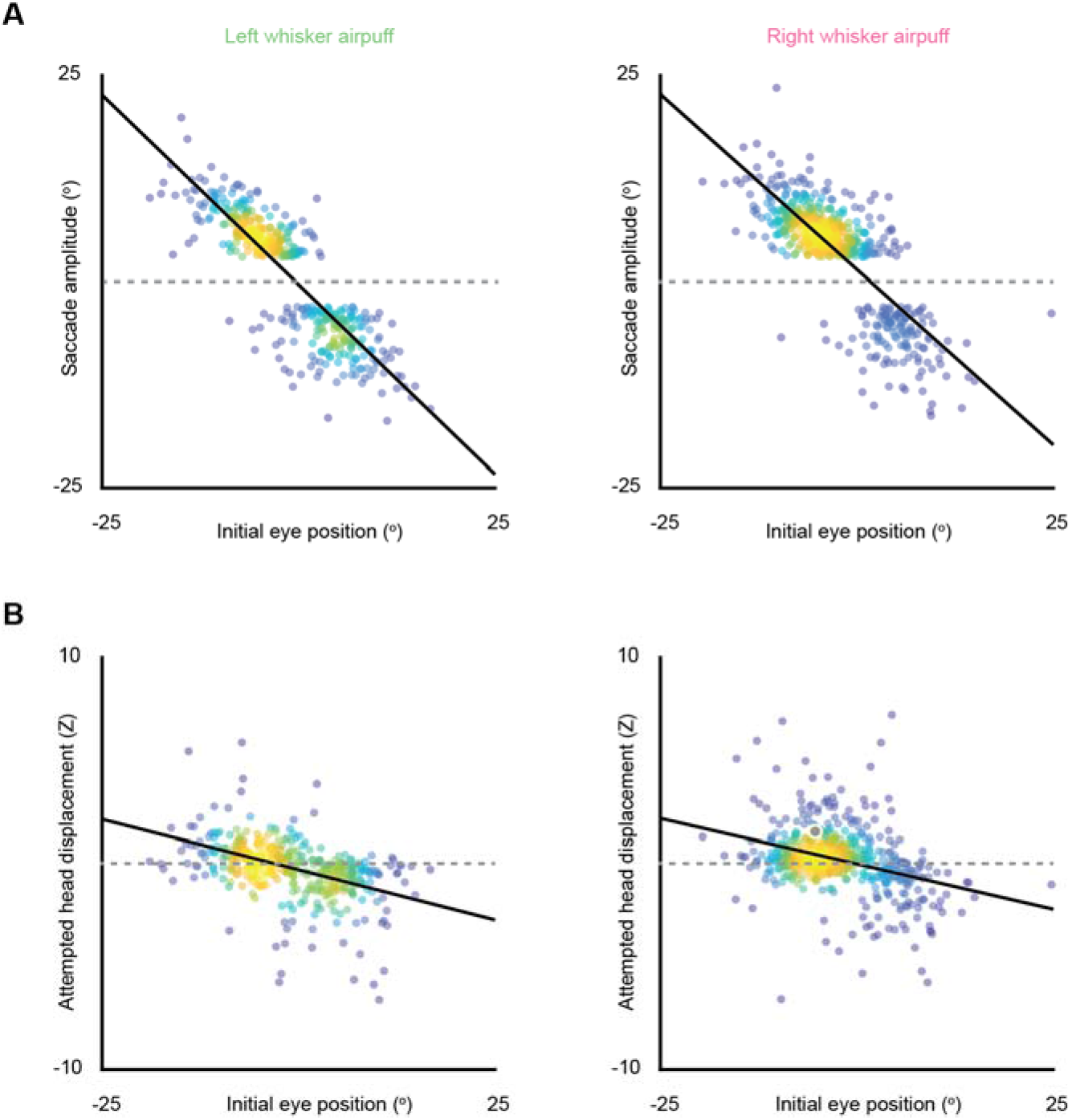
Whisker-evoked saccade and attempted head movement direction and amplitude vary according to initial eye position. **(A)** Saccade amplitude as a function of initial eye position for left and right whisker airpuff-evoked gaze shift. Each dot corresponds to a single saccade. Brighter areas indicate higher densities of points. Linear fits for left and right airpuffs, respectively: slope = −0.92 and −0.85, R^2^ = 0.74 and 0.63, p < 10^-10^ and p < 10^-10^, n = 441 and 606. **(B)** Attempted head movement amplitude as a function of initial eye position for left and right whisker airpuff-evoked gaze shifts. Linear fits for left and right airpuffs, respectively: slope = −0.10 and −0.09, R^2^ = 0.19 and 0.13, p < 10^-10^ and p < 10^-10^, n = 441 and 606. Dashed lines correspond to 0. Values above and below dashed line correspond to attempted rightward and leftward movements, respectively.

**Supplementary Figure 9.**
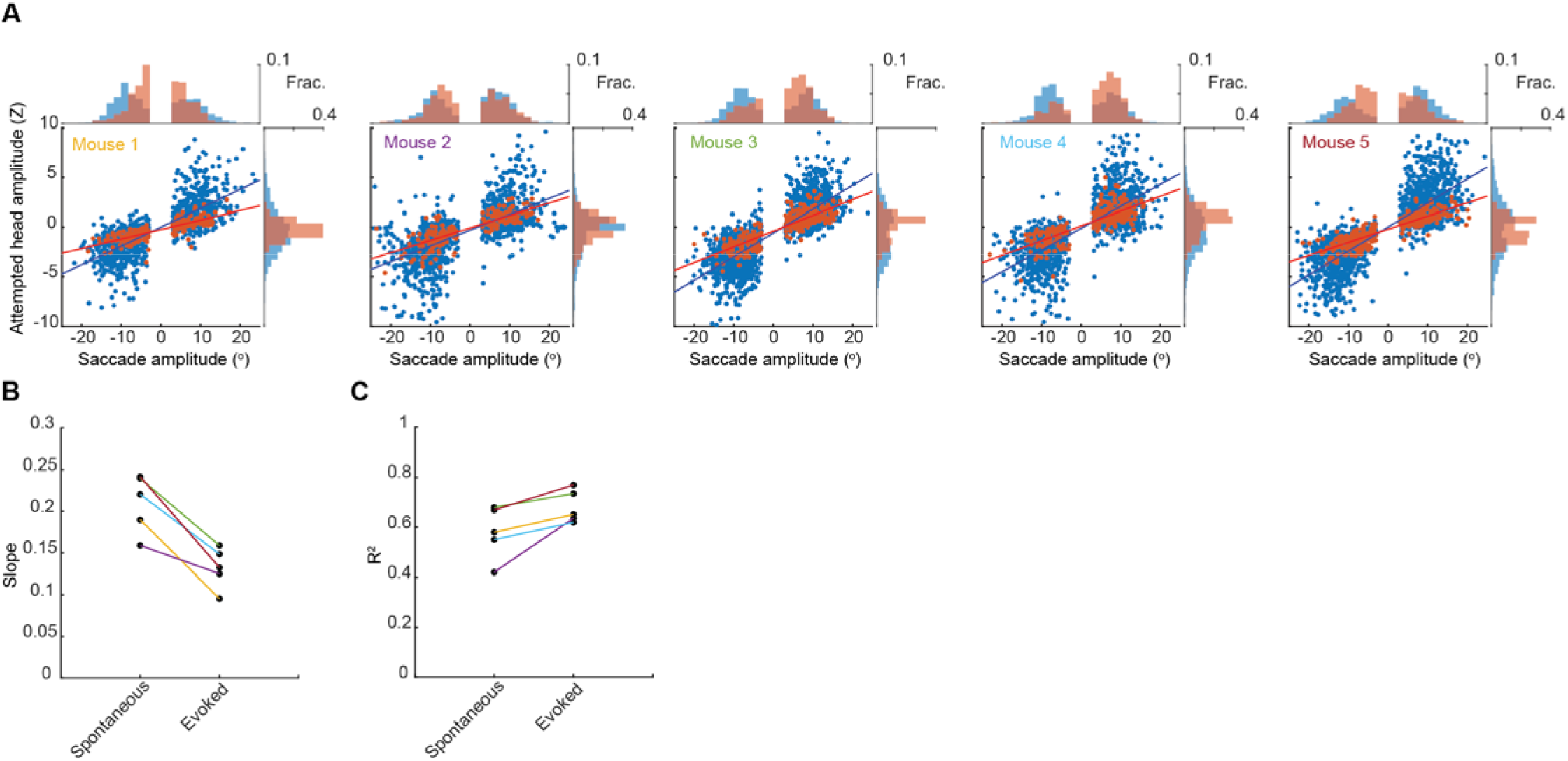
Gain of head-eye coupling variability across mice. (A) Head-eye amplitude coupling of spontaneous (blue) and ear airpuff-evoked saccades (red) for five individual mice. Each dot corresponds to a single gaze shift. The spontaneous and evoked regression slopes were significantly different for all mice (p < 10^-5^, permutation test). Histograms above and beside scatter plot indicate distributions of saccade and attempted head movement amplitudes.**(B, C)** Slope and R^2^ for linear fits to spontaneous and evoked gaze shifts for each mouse. Colored lines indicate values for the different mice in A.

**Supplementary Figure 10.**
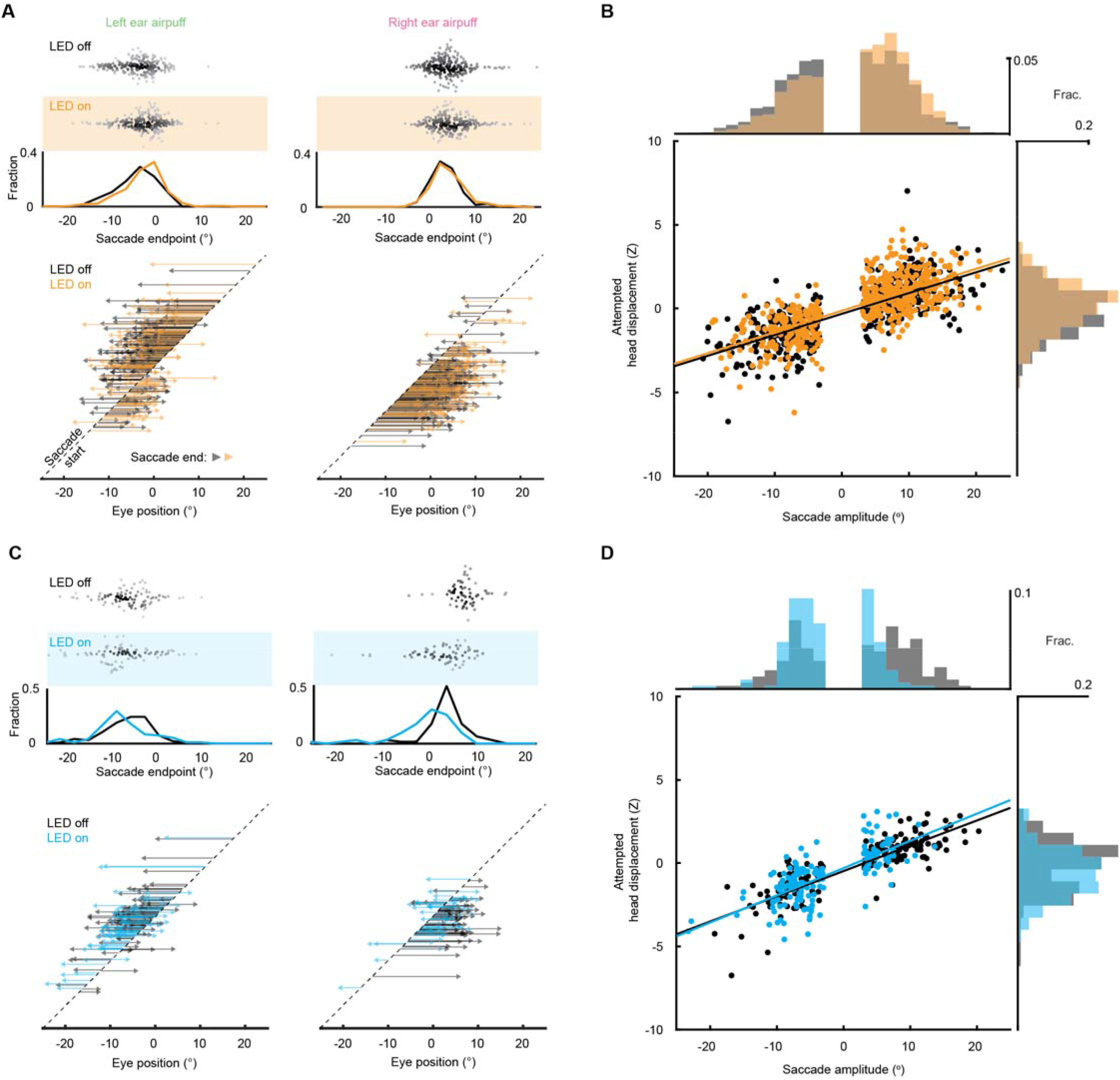
Controlling for the effects of initial eye position on superior colliculus manipulations. **(A)** Effects of SC optogenetic inhibition on saccade endpoints for trials matched for initial eye position. Top, scatter plots and histograms of endpoints for control (white background) and LED on (orange background) trials. Middle, endpoint histograms for control (black) and LED on (orange) trials. Bottom, saccade vectors for control (black) and LED on (orange) trials. **(B)** Head-eye amplitude coupling during ear airpuff-evoked gaze shifts for control (black) and LED on (orange) trials matched for initial eye position. Each dot represents an individual gaze shift. Control: R^2^ = 0.57, slope = 0.124, p < 10^-10^. LED on: R^2^ = 0.53, slope = 0.125, p < 10^-10^. Control and LED-on regression slopes were not significantly different (p = 0.41, permutation test) Histograms above and beside scatter plot show distributions of saccade amplitude and head displacement, respectively. Distribution means were significantly different (p = 0.002 for saccades, p < 10^-5^ for attempted head movements, permutation test). **(C)** Effects of weak SC optogenetic stimulation on saccade endpoints for trials matched for initial eye position. Top, scatter plots and histograms of endpoints for control (white background) and LED on (blue background) trials. Middle, histograms of endpoints for control (black) and LED on (blue) trials. Bottom, saccade vectors for control (black) and LED on (blue) trials. **(D)** Head-eye amplitude coupling during ear airpuff-evoked gaze shifts for control (black) and LED on (blue) trials matched for initial eye position. Each dot represents an individual gaze shift. Control: R^2^ = 0.74, slope = 0.15, p < 10^-10^. LED on: R^2^ = 0.52, slope = 0.164, p < 10^-10^. Control and LED-on regression slopes were not significantly different (p = 0.35, permutation test) Histograms above and beside scatter plot show distributions of saccade amplitude and head displacement, respectively. Means were significantly different (p < 10^-5^ for saccades, p = 0.002 for attempted head movements, permutation test).

## Materials and Methods

### Mice

All experiments were performed according to Institutional Animal Care and Use Committee standard procedures. C57BL/6J wild-type (Jackson Laboratory, stock 000664) mice between 2 and 6 months of age were used. Mice were housed in a vivarium with a reversed 12:12 h light:dark cycle and tested during the dark phase. No statistical methods were used to predetermine sample size. Behavioral experiments were not performed blinded as the experimental setup and analyses are automated.

### Surgical procedures

Mice were administered carprofen (5 mg/kg) 30 minutes prior to surgery. Anesthesia was induced with inhalation of 2.5% isoflurane and buprenorphine (1.5 mg/kg) was administered at the onset of the procedure. Isoflurane (0.5-2.5% in oxygen, 1 L/min) was used to maintain anesthesia and adjusted based on the mouse’s breath and reflexes. For all surgical procedures, the skin was removed from the top of the head and a custom titanium headplate was cemented to the leveled skull (Metabond, Parkell) and further secured with dental cement (Ortho-Jet powder, Lang Dental). Craniotomies were made using a 0.5 mm burr and viral vectors were delivered using pulled glass pipettes coupled to a microsyringe pump (Micro4, World Precision Instruments) on a stereotaxic frame (Model 940, Kopf Instruments). Following surgery, mice were allowed to recover in their home cages for at least 1 week.

### Viral injections and implants

Coordinates for SC injections were ML: 1.25 mm, AP: 0.7 mm (relative to lambda), DV: −1.9 and −2.1 mm (100 nL/depth). Coordinates for SC implants were ML: 1.25 mm, AP: 0.7 mm (relative to lambda), DV: −2.0 mm. Fiber optic cannulae were constructed from ceramic ferrules (CFLC440-10, Thorlabs) and optical fiber (400 mm core, 0.39 NA, FT400UMT) using low-autofluorescence epoxy (F112, Eccobond).

### Behavioral procedures

To characterize stimulus-evoked gaze shifts (figures 1, 2, 3 and related supplements), data were collected from 5 mice over 53 days (maximum of 1 session/mouse/day). Session types were randomly interleaved to yield a total of 6 ear airpuff sessions, 6 ear tactile sessions, 6 whisker airpuff sessions, 10 auditory airpuff sessions, and 5 visual sessions. During experiments, headplated mice were secured in a custom 3D-printed mouse holder. Timing and synchronization of the behavior were controlled by a microcontroller (Arduino MEGA 2560 Rev3, Arduino) receiving serial commands from custom Matlab scripts. All behavioral and data acquisition timing information was recorded by a NI DAQ (USB-6001) for post hoc alignment. All experiments were performed using awake mice. Left and right stimuli were randomly selected and presented at intervals drawn from a 7-12 s uniform distribution. Each session consisted of 350 stimulus presentations and lasted ∼55 minutes. No training or habituation was necessary.

### Stimuli

Airpuff stimuli were generated using custom 3D-printed airpuff nozzles (1.5 mm wide, 10 mm long) connected to compressed air that was gated by a solenoid. 3D-printed nozzles were used to standardize stimulus alignment across experimental setups but similar results were obtained in preliminary experiments using a diverse array of nozzle designs. For whisker airpuffs, the nozzles were spaced 24 mm apart and centered 10 mm beneath the mouse’s left and right whiskers. For ear airpuffs, the nozzles were directed toward the ears while maintaining 10 mm of separation between the nozzles and the mouse. For auditory-only airpuffs, the nozzles were directed away from the mouse while maintaining the same azimuthal position as the ear airpuffs. For tactile-only stimulation, the ears were deflected using a thin metal bar coated in epoxy to soften its edges (7122A37, McMaster). A stepper motor (Trinamic, QSH2818-32-07-006 and TMC2208) was programmed to sweep the bar downward against the ear before sweeping back up. The stepper motor was sandwiched between rubber pads (8514K61, McMaster) and elevated on rubber pedestals (20125K73, McMaster) to reduce any sound due to vibration. For visual stimulation, white LEDs (COM-00531, Sparkfun) were mounted 6 inches from the mouse at the same azimuthal position as the airpuff nozzles.

### Eye tracking

The movements of both left and right eyes was monitored at 100 Hz using two high-speed cameras (BFS-U3-28S5M-C, Flir) coupled to a 110 mm working distance 0.5X telecentric lens (#67-303, Edmund Optics). A bandpass filter (FB850-40, Thorlabs) was attached to the lens to block visible illumination. Three IR LEDs (475-1200-ND, DigiKey) were used to illuminate the eye and one was aligned to the camera’s vertical axis to generate a corneal reflection. Videos were processed post hoc using DeepLabCut, a machine learning package for tracking pose with user-defined body parts (Mathis et al., 2018). Data in this paper were analyzed using a network trained on 1000 frames of recorded behavior from 8 mice (125 frames per mouse). The network was trained to detect the left and right edges of the pupil and the left and right edges of the corneal reflection. Frames with a DeepLabCut-calculated likelihood of *P* < 0.90 were discarded from analyses. Angular eye position (E) was determined using a previously described method developed for C57BL/6J mice (Sakatani and Isa, 2004).

### Attempted head rotation tracking

Attempted head rotations were measured using a 3D-printed custom headplate holder coupled to a load cell force sensor (Sparkfun, SEN-14727). Load cell measurements (sampling frequency 80 Hz) were converted to analog signals and recorded using a NI DAQ (sampling frequency 2000 Hz). The data were then low-pass filtered at 80 Hz using a zero-phase second-order Butterworth filter and then downsampled to match the pupil sampling rate.

### Optogenetics

Optogenetic experiments were performed using the ear airpuff nozzles. Fiber optic cables were coupled to implanted fibers and the junction was shielded with black heat shrink. A 470 nm fiber-coupled LED (M470F3, Thorlabs) was used to excite ChR2-expressing neurons, and a 545 nm fiber-coupled LED (UHP-T-SR, Prizmatix) was used to inhibit eNpHR3.0-expressing neurons. Optogenetic excitation during WISLR sessions was delivered on a random 50% of trials using 1 s of illumination centered around airpuff onset.

#### SC inhibition

For optogenetic inactivation of SC neurons, AAV1.hSyn.eNpHR3.0 was injected into the right SC of 5 wild-type mice (0.6 AP, 1.1 ML, −2.1 and −1.9 DV; 100 nl per depth). Experiments were performed 35-40 days post injection. LED power was 12 mW. Mice underwent 5 sessions each.

#### SC stimulation

For subthreshold optogenetic stimulation of SC neurons, AAV1.CaMKIIa.hChR2(H134R)-EYFP was injected into the right SC of 4 wild-type mice. Experiments were performed 67-71 days post injection. LED power was individually set to an intensity that did not consistently evoke saccades upon LED onset (50-120uW). Mice underwent 5 sessions each.

### Histology

For histological confirmation of fiber placement and injection site, mice were perfused with PBS followed by 4% PFA. Brains were removed and post-fixed overnight in 4% PFA, and stored in 20% sucrose solution for at least 1 day. Brains were sectioned at 50 μm thickness using a cryostat (NX70, Cryostar), every third section was mounted, and slides were cover-slipped using DAPI mounting medium (Southern Biotech). Tile scans were acquired using a confocal microscope (LSM700, Zeiss) coupled to a 10X air objective.

### Saccade detection

Saccades were defined as eye movements that exceeded 100°/s, were at least 3° in amplitude, and were not preceded by a saccade in the previous 100 ms. The initial positions and endpoints of saccades were defined as the first points at which saccade velocity rose above 30°/s and fell below 20°/s, respectively. Analyses focused on horizontal saccades because saccades were strongly confined to the azimuthal axis (Supplementary Fig. 1).

### Behavioral analysis

Eye position analyses were performed using the averaged left and right pupil positions, and the mean eye position was subtracted from each session prior to combining data across sessions and mice. Similarly, attempted head rotation data was Z-scored prior to combining data across sessions and mice. To quantify stimulus-evoked saccade probability, we calculated the fraction of trials in which a saccade occurred in the 100 ms period following stimulus onset (i.e. the response window). To quantify stimulus-evoked attempted head rotation probability in Figure 1, we calculated the fraction of trials in which the Z-scored head sensor reading exceeded 0.25Z 150 ms following stimulus onset. To determine the baseline head movement probability, we calculated the fraction of trials in which the head sensor reading exceeded 0.25Z between −500 ms and −350 ms relative to saccade onset.

To examine saccade endpoints, we first identified trials in which mice maintained fixation in the 500 ms preceding saccade onset. We then considered stimulus-evoked saccades those occuring within 100 ms of stimulus onset.

To examine attempted head movement latencies, we first identified trials in which the mice maintained head fixation from −1 to −0.5 s prior to saccade onset and used this period as the baseline. Latency was defined as the first frame between −0.5 and 0.5 s relative to saccade onset when the attempted head movement amplitude exceeded 5 standard deviations from that trial’s baseline.

To examine head-eye amplitude coupling during spontaneous and stimulus-evoked gaze shifts, we identified trials in which the mouse maintained fixation in the 500 ms preceding saccade onset. We defined attempted head rotation amplitude as the load cell value 150 ms following saccade onset (the time point at which average load cell value plateaus during stimulus-evoked gaze shifts (Fig. 3)). For certain analyses, we identified saccades matched (without replacement) for initial eye position and/or saccade amplitude using Euclidean distance as a metric and a 3° distance cutoff.

Tests for statistical significance are described in the text and figure legends. Data were shuffled 10,000 times to generate a null distribution for permutation tests.

